# Resolving the Functional Significance of *BRCA1* RING Domain Missense Substitutions

**DOI:** 10.1101/092619

**Authors:** Andrew Paquette, Kayoko Tao, Kathleen A Clark, Alex W Stark, Judith Rosenthal, Angela K Snow, Russell Bell, Bryony A Thompson, Joshua Unger, Brett A Milash, Lisa Pappas, Jason Gertz, Katherine E Varley, Alun Thomas, Ken Boucher, William D Foulkes, David E Goldgar, Sean V Tavtigian

## Abstract

**Part 1:** Development and calibration of suitably accurate functional assays for *BRCA1* RING domain and BRCT domain missense substitutions could dramatically accelerate clinical classification of rare missense substitutions observed in that gene. Leveraging data from 68,000 full sequence tests of *BRCA1* and *BRCA2*, plus data from the limited number of already classified *BRCA1* RING domain missense substitutions, we used logistic regression and related techniques to evaluate three *BRCA1* RING domain assays. These were recently described high throughput yeast 2-hybrid and E3 ubiquitin ligase assays, plus a newly developed mammalian 2-hybrid assay. While there were concerns about the accuracy of the yeast 2-hybrid assay and the indirect nature of the ubiquitin ligase assay, the mammalian 2-hybrid assay had excellent correlation with existing missense substitution classifications. After calibration, this assay contributed to classification of one newly reported *BRCA1* missense substitution. In principal, the mammalian 2-hybrid assay could be converted to a high-throughput format that would likely retain suitable accuracy.

**Part 2:** How does one achieve clinically applicable classification of the vast majority of all possible sequence variants in disease susceptibility genes? BRCA1 is a high-risk susceptibility gene for breast and ovarian cancer. Pathogenic protein truncating variants are scattered across the open reading frame, but all known missense substitutions that are pathogenic because of missense dysfunction are located in either the amino-terminal RING domain or the carboxy-terminal BRCT domain. Heterodimerization of the BRCA1 and BARD1 RING domains is a molecularly defined obligate activity. Hence, we tested every BRCA1 RING domain missense substitution that can be created by a single nucleotide change for heterodimerization with BARD1 in a Mammalian 2-hybrid (M2H) assay. Downstream of the M2H laboratory assay, we addressed three additional challenges: assay calibration, validation thereof, and integration of the calibrated results with other available data such as computational evidence and patient/population observational data to achieve clinically applicable classification. Overall, we found that about 20% of BRCA1 RING domain missense substitutions are pathogenic. Using a Bayesian point system for data integration and variant classification, we achieved clinical classification of about 89% of observed missense substitutions. Moreover, among missense substitutions not present in the human observational data used here, we find an additional 47 with concordant computational and functional assay evidence in favor of pathogenicity; these are particularly likely to be classified as Likely Pathogenic once human observational data become available.

## INTRODUCTION to PART 1

*BRCA1* (MIM #113705) and its heterodimerization partner *BARD1* (MIM #601593) share two highly conserved domains: an N-terminal RING domain and a pair of C-terminal BRCT repeats. Even though the RING and BRCT domains comprise only about 17% of the length of *BRCA1*, all of the missense substitutions in this protein that are known to be pathogenic occur in one of these two domains (http://hci-exlovd.hci.utah.edu/home.php?select_db=BRCA1), unless the underlying nucleotide change is spliceogenic. While severely dysfunctional missense substitutions in either of these domains are associated with increased cancer risk (reviewed in Clark et al., 2012), it is still unclear how non-spliceogenic RING missense substitutions elicit their pathogenicity

In contrast with protein truncating variants, it is often difficult to know what effect, if any, missense substitutions will have on protein function. Over the last 12 years, we and others have developed a Bayesian “integrated evaluation” or “multifactorial likelihood model” for evaluation of Variants of Unclear Significance (VUS) in *BRCA1* and *BRCA2* (Goldgar et al. 2004, 2008; Easton et al. 2007). This integrated evaluation combines a sequence analysis-based prior probability of pathogenicity (Prior_P) (Tavtigian et al., 2008; Vallée et al., 2016) with observational data from the patient and/ or tumor, expressed as odds in favor of pathogenicity (Odds_Path) to arrive at a posterior probability of pathogenicity (Post_P) (Lindor et al. 2012; Vallée et al. 2012; Thompson et al., bioRxiv 079418; doi: http://dx.doi.org/10.1101/079418). The resulting posterior probability is then converted to one of five qualitative classes, based on cutpoints considered to be clinically relevant (Plon et al. 2008). Using this quantitative approach, 155 missense substitutions have now been classified in *BRCA1* (http://hci-exlovd.hci.utah.edu/home.php?select_db=BRCA1). Of these, ten are located in the RING domain, with eight falling into one of the two pathogenic classes (IARC Class 4 or 5) and two falling into one of the two neutral classes (IARC Class 1 or 2).

Two well recognized activities of BRCA1 reside in its first 300 amino acids. The interval from roughly Ala4 to Ala102 encodes a C3HC4 RING finger and a set of helical bundles that together enable heterodimerization with the homologous domain of BARD1 (Brzovic et al., 2001; L. Wu et al., 1996). In addition, the interval from roughly Ala4 to Asp300 encodes E3 ubiquitin ligase activity (Nishikawa et al. 2004). The BRCA1:BARD1 interaction influences the abundance and stability of BRCA1 (Wu et al. 2010), is necessary for early recruitment of BRCA1 to sites of DNA double strand breaks (Li and Yu, 2013), and fully activates the BRCA1 E3 ubiquitin ligase activity (Baer and Ludwig, 2002; Brzovic et al., 2003; Hashizume et al., 2001). To our knowledge, there are no well-documented *BRCA1* separation of function mutations that severely damage the BARD1 interaction without also dramatically reducing E3 ubiquitin ligase activity. On the other hand, the *BRCA1* missense substitution p.Ile26Ala, which does not dramatically reduce BARD1 interaction, specifically disrupts the interaction between BRCA1 and its cognate E2 ligases, selectively inhibiting E3 ligase activity by blocking the transfer of ubiquitin to substrates bound to BRCA1 (Brzovic et al., 2003; Wu et al., 2008). Interestingly, mice homozygous for this RING domain separation of function missense substitution are no more tumor prone than their wild-type littermates (Shakya et al. 2011). In contrast, RING variants that disrupt heterodimer formation result in DNA repair defects and loss of tumor suppression (Ransburgh et al., 2010). Nonetheless, Starita et al reported that BRCA1:BARD1 heterodimer formation alone is a poor predictor of the pathogenicity of RING missense substitutions, implying that the E3 ligase activity does make an important contribution (Starita et al. 2015).

Previously, we used data from 68,000 full sequence tests of *BRCA1* and *BRCA2* performed at Myriad Genetics to develop and calibrate computational algorithms for evaluation of missense substitution severity and splice variant severity in these two genes (Tavtigian et al. 2008; Vallée et al. 2016). Here, linking the Myriad *BRCA1* test data to the nearly comprehensive Yeast 2-hybrid (Y2H) BRCA1:BARD1 interaction assay data and phage display BRCA1 E3 ligase activity data published recently by Starita et al. (2015), we test two 1-sided hypotheses: (1) as measured by the Y2H assay, that loss of BRCA1:BARD1 interaction is predictive of pathogenicity, and (2) as measured by the phage-display E3 ligase assay, that loss of BRCA1 auto-ubiquitination is predictive of pathogenicity. Guided by the results from these hypothesis tests, we develop and calibrate an accurate functional assay, results from which can be included within the Bayesian integrated evaluation framework to assess the pathogenicity of *BRCA1* RING missense substitutions. We then report progress towards conversion of that assay to a high-throughput format that could evaluate all possible *BRCA1* amino terminus substitutions.

## METHODS

### MIM # and accession #

*BRCA1* is MIM# 113705, and exonic variant coding used here is based on NM_007294.3.

*BARD1* is MIM# 601593, and exonic variant coding used here is based on NM_000465.3.

### Dataset

The dataset comprised results of full sequence tests carried out at Myriad Genetic Laboratories, as used previously in Easton et al. (2007) and Tavtigian et al. (2008) for modelling of risk associated with *BRCA1/2* sequence variation. The analyses described here are based on results of full sequence tests of both genes from 68,000 BRACAnalysis subjects of whom 4,867 were reported to carry a pathogenic *BRCA1* variant and 3,561 were reported to carry a pathogenic *BRCA2* variant. For a test to have been performed, a test request form must have been completed by the ordering health care provider, and the form must have been signed by an appropriate individual indicating that “informed consent has been signed and is on file”. The mutation screening data are arranged by sequence variant rather than by subject. The dataset includes nucleotide and amino acid nomenclature specifications for all of the exonic single nucleotide substitutions – silent, missense, or nonsense – observed from the 68,000 patient mutations screening set; these are all of the observational data required to calculate the Enrichment Ratio for Single Nucleotide Substitutions (ERS) (Tavtigian et al. 2008).

Analyses of the personal and family history of tested probands to calculate family history likelihood ratios (FamHx-LRs) derive from a virtually identical series of subjects used previously (Easton et al. 2007). However, this dataset also includes frameshifts, in-frame indels, and sequence variants falling in the intronic portions of the splice junction consensus regions from - 20 to +6 of the protein coding exons. We refer to these two overlapping data sets as the B1&2 68K set.

### Additional Subjects

Independent to the B1&2 68K set, a family with the rare missense substitution *BRCA1* p.P34S was identified. Informed consent was obtained before subjects provided a cancer personal & family history, and were tested for carriage of this variant.

### Enrichment Ratio for Single Nucleotide Substitutions (ERS) calculations

The ERS is similar in spirit to the traditional population genetics measure d_N_/d_S_ (d_N_ is the non-synonymous substitution rate and d_S_ is the synonymous substitution rate per site), where a d_N_/d_S_ ratio greater than 1.0 is indicative of positive selection (Yang 1998). For each nucleotide in a canonical DNA sequence, there are three possible single nucleotide substitutions. However, these substitutions are not equally likely to occur because of differences in the underlying substitution rate constants. Using the dinucleotide substitution rate constants given by Lunter and Hein (2004), averaging sense and antisense orientations, we can estimate a relative substitution rate for every possible single-nucleotide substitution to a DNA sequence, **r_i_**. The probability that a new sequence variant (i.e., a new germline sequence variant at the moment that it comes into existence) will fall into a particular algorithmically defined class **c**† is given by the ratio of the sum of the relative substitution rates of the variants belonging to the class **c** divided by the sum of all relative substitution rates:

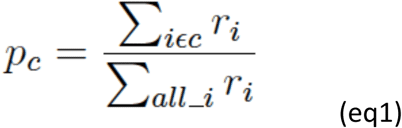

Hence, under the null hypothesis of no selection, we can obtain from the total number of variants observed in a mutation screening study, **o_T_**, the number expected in any class, **e_C_=p_C_o_T_**, and compare this to the actual number observed, **o_C_**. Thus, in general, we define the ERS for any class of substitutions **c** as the observed / expected ratio for that class normalized by the same ratio for silent (i.e., synonymous) substitutions but excluding the few silent substitutions that are likely to be spliceogenic:

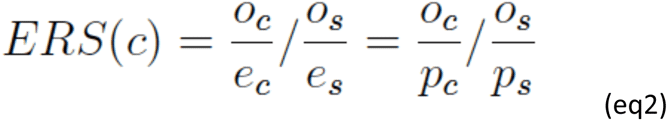

† For this discussion, an “algorithmically defined class” of variants is a class of variants that can be unambiguously specified by an algorithm. One example could be, given a specified protein multiple sequence alignment, all substitutions that fall at an invariant position in the alignment and have a Grantham Score ≥65. Another could be, given a functional assay that evaluated essentially all possible substitutions in a given protein domain, all substitutions resulting in <60% of wild-type (wt) activity in the functional assay.

### Regressions of high-throughput yeast 2-hybrid and phage display ubiquitin E3 ligase assay results against B1&2 68K data

To test the hypothesis that the Y2H assay is a predictor of pathogenicity for missense substitutions in the BARD1 interaction domain of *BRCA1*, we performed a logistic regression of the presence of the variant (yes/ no) in the B1&2 68K data set as a function of the result of the Starita et al. (2015) Y2H assay, controlling for the nucleotide substitution rate constant of each substitution. This analysis was limited to *BRCA1* amino acid positions 2-103. Because raw Y2H assay results were skewed towards 100% activity whereas exponentiated Y2H assay results gave a more symmetric distribution, exponentiated Y2H results were used in the regression. Because the log of the substitution rate constants was closer to a Gaussian distribution than the raw rate constants, log rate was used in the regression.

To test the hypothesis that the E3 ubiquitin ligase assay is a predictor of pathogenicity for missense substitutions in the E3 ligase domain, but outside of the BARD1 binding domain, we performed a logistic regression of the presence of the variant (yes/ no) in the B1&2 68K data set as a function of the result of the Starita et al. (2015) E3 ligase assay, controlling for the nucleotide substitution rate constant of each substitution. This analysis was limited to *BRCA1* amino acid positions 104-300. The distribution of raw E3 ligase assay results was adequately symmetric for use in the logistic regressions as raw data.

### 1-by-1 mammalian 2-hybrid assay

#### Cell line development

The firefly luciferase reporter pGL4.31 (Promega), was re-engineered for puromycin resistance, and then stably incorporated into HEK293 cells (ATCC CRL-1573) using the PiggyBac transposon system (System Biosciences). Chromatin insulators and PiggyBac terminal repeats were added to each end of the reporter to block transgene silencing and facilitate transposition, respectively. These sequences were PCR amplified from PB531A-2 (System Biosciences).

500,000 HEK293 cells/well were plated in a 6-well plate 18-24 hours before transfection. 500 ng of PiggyBac pGL4.31 Puro and 200 ng of PiggyBac transposase (PB200A-1, System Biosciences) were co-transfected with 8 μl of FugeneHD (Promega). The cells were then subjected to one week of selection (2 μg/ml puromycin), at which point individual clones were transferred to 96-well plates via FACS. Luciferase activity was characterized by the 2-hybrid system described below, using known neutral *BRCA1* RING missense substitutions and wt *BARD1*. The clone with the highest relative luminescence signal (∼1×10^6^ RLU) was used to generate the stable cell line (HEK293 PiggyBac pGL4.31 Puro) used for subsequent mammalian 2-hybrid assays.

#### Cell line maintenance

HEK293 PiggyBac pGL4.31 Puro cells were grown in high-glucose DMEM (Gibco) supplemented with 10% fetal bovine serum (Gibco), Na-pyruvate (Gibco, 110 mg/L), penicillin/streptomycin (Gibco, 5,000 U/ml), and puromycin (Gibco, 2 μg/ml). Cells were passaged with phenol red-free TrypLE (Gibco, 1x) as needed at a 1:10 split.

#### Plasmid construction

Mammalian 2-hybrid assay vectors, pACT (E246A, Promega) and pBIND (E245A, Promega), were modified by moving VP16 and GAL4 to the C-terminus. The first 184 amino acids of *BRCA1*, fused to 3x FLAG, was cloned into pBIND immediately upstream of GAL4, while the first 200 amino acids of *BARD1*, fused to HA, was cloned into pACT immediately upstream of VP16. *BRCA1* and *BARD1* cDNA templates were obtained from pCL-MFG-BRCA1 (Addgene plasmid #12341) (Ruffner and Verma 1997) and BARD1 pET28a (Addgene plasmid #12646) (Brzovic et al., 2006), respectively. Linker sequences were placed between all coding regions to limit steric hindrance.

#### Mammalian 2-hybrid assay

BRCA1:BARD1 heterodimer formation was evaluated by co-transfecting HEK293 PiggyBac pGL4.31 Puro cells with wt *BARD1* (pACT_BARD1 1-200), and either wt *BRCA1* or various *BRCA1* RING missense substitutions (pBIND_BRCA1 1-184). Three independent clones of each *BRCA1* RING missense substitution were individually incorporated into the pBIND vector using the QuikChange lightning site-directed mutagenesis kit (Agilent). The presence of each variant, and lack of additional mutations in the *BRCA1* coding sequence, was confirmed by Sanger sequencing (data not shown).

HEK293 PiggyBac pGL4.31 Puro cells were transfected using TransIT-293 (Mirus). Briefly, 13,000 cells/well were plated on a 96-well microplate in antibiotic-free, phenol-free RPMI (Gibco) supplemented with 10% FBS. 18-24 hours later, 50 ng of wt *BARD1* (pACT_BARD1 1-200) and 50 ng of either wt *BRCA1* (pBIND_BRCA1 1-184), an empty vector (pBIND) or various *BRCA1* RING missense substitutions were co-transfected at a 1:1 molar ratio in 10 μl of phenol-free Opti-MEM (Gibco), with TransIT-293 being used at a 3:1 ratio (0.4 μl/well). Each individual *BRCA1* missense clone was assayed in triplicate on a single day (batch). The three independent clones of a given missense substitution were assayed in different batches. Wt *BARD1* was replaced with an empty vector on the right half of each plate as a background control. Firefly and Renilla luciferase expression were quantified 48 hours later on a Glomax 96 Microplate Luminometer using the Dual-Glo Luciferase Assay System (Promega). Observed Firefly luciferase activity was normalized for transfection efficiency by dividing Firefly by Renilla, and multiplying by 1,000. This normalized activity (measured in triplicate for each clone) was then converted to % wt activity by dividing each normalized measure by the average activity of the wt control on the same plate. This value was then multiplied by the inverse variance of each clone (weight), summed, and divided by sum weight of all three clones to arrive at a weighted average of % wt activity. A standard 95% CI is reported for each *BRCA1* RING missense substitution.

### Co-Immunoprecipitation

HEK293 cells were transfected using Polyethylenimine (PEI) (Polysciences Incorporated). Briefly, 3×10^6^ HEK293 cells were plated on a 10cm^2^ plate in high-glucose DMEM (Gibco) supplemented with 10% fetal bovine serum (Gibco), Na-pyruvate (Gibco, 110 mg/L), and penicillin/streptomycin (Gibco, 5,000 U/ml). 18-24 hours later, 2.5 μg of wt *BARD1* (pACT_BARD1 1-200) or an empty vector (EV), and either wt *BRCA1* or various *BRCA1* RING missense substitutions (pBIND_BRCA1 1-184) were co-transfected with 500 ng of pBIG (GFP control plasmid), with PEI (1 mg/ml) being used at a 2:1 ratio.

Cell lysates were harvested in phospho-protecting lysis buffer (PPLB) 48 hours post-transfection and immunoprecipitation performed from clarified cell lysates using α-FLAG (M2) antibody and Protein G Sepharose beads (Engel et al., 2010). Immune complexes and clarified lysates from each transfection were probed using mouse monoclonal α-FLAG (M2) and α-tubulin (B-5-1-2) from Sigma Aldrich, rabbit polyclonal HA (ab9119) from Abcam, rabbit polyclonal GFP (sc-8334) from Santa Cruz Biotechnology, IRDye 800CW Goat anti-Mouse (925-32210), and IRDye 680RD Goat anti-Rabbit (925-68071) from LI-COR Biosciences.

To avoid IgG heavy and light chain interference, immune complexes were denatured in Bolt LDS sample buffer without dithiothreitol (DTT), and heated to 70°C for 10 minutes. Clarified cell lysates were denatured in a similar manner, with the exception that the sample buffer was supplemented with 50 mM DTT. Proteins were fractionated by electrophoresis on 8% Bis-Tris Plus gels (Invitrogen) in 1X Bolt MES SDS Running Buffer (Novex) for 30 minutes at 165 Volts. Transfers were conducted on iBlot 2 PVDF transfer stacks (Invitrogen) using pre-programmed template P0.

### High-throughput mammalian 2-hybrid assay

To explore conversion of the existing 1-by-1 mammalian 2-hybrid assay into a high-throughput screen, the 2-hybrid luciferase reporter plasmid was converted to two independent fluorescent reporters, one with ZsGreen and blasticidin resistance, and the other with Tdtomato and puromycin resistance. These two fluorescent reporters were then targeted with homology arms to the common integration site of the non-pathogenic adeno-associated virus (AAVS1) and ROSA26, respectively. For this transfection, 1 μg of each CRISPR and donor plasmid was co-transfected with 24 μl of PEI (1 mg/ml). After co-transfection into HEK293 cells with the appropriate CRISPR/cas9 reagents and double selection with puromycin (2 μg/ml) and blasticidin (Gibco, 15 μg/ml), a pool of cells was harvested for downstream experiments. PCR confirmed that the pool included cells with stably integrated ZsGreen and Tdtomato. In some of these cells, Tdtomato was located at ROSA26, but it was not possible to confirm location of ZsGreen at AAVS1.

BRCA1:BARD1 heterodimer formation was evaluated as described in the 1-by-1 mammalian 2-hybrid assay, with the following modifications: 625,000 ZsGreen/Tdtomato cells/well were plated in a 6-well plate 18-24 hours before transfection. 525 ng of pBIND_BRCA1 1-184 and 469 ng of pACT_BARD1 1-200 were co-transfected at a 1:1 molar ratio, and PEI (1 mg/ml) was used at a 6:1 ratio. 48 hours post-transfection, the six wells of cells were mixed and FACS sorted into four bins based on Tdtomato expression. Each of these bins was further sorted into four bins based on ZsGreen expression. RNA was isolated from each bin along the dual-expression main diagonal using the Direct-zol RNA miniprep kit (Zymo Research), and converted to cDNA using the SuperScript III first-strand synthesis system (Invitrogen). The RING sequence of *BRCA1* was then RT-PCRd with an intron-spanning primer and sequencing libraries prepared using the Ovation Ultralow Library System (NUGEN # 0329), excluding the end-repair step and using a different barcode for each of the four main diagonal red-green fluorescence intensity bins. The libraries were sequenced on a Illumina MiSeq channel using the MiSeq 300 bp Cycle Paired-end sequencing protocol.

### Statistical analyses

Regressions of high-throughput Y2H and phage display ubiquitin E3 ligase assay results against B1&2 68K data were performed in R version 3.2.1 (R Core Team, 2015). Regressions to calibrate the Mammalian 2-hybrid assay results were performed in Stata 11.0 (StataCorp).

## RESULTS

### Re-analysis of Y2H and E3 ubiquitin ligase assay data

Because *BRCA1* is a breast/ ovarian cancer susceptibility gene, BRACAnalysis is a sequencing test fundamentally designed to detect pathogenic sequence variants in *BRCA1* (and *BRCA2*), and the B1&2 68K set of subjects was strongly enriched in individuals with a personal and/or family history of breast/ ovarian cancer, we expect to observe a disproportionate excess of sequence variants in the B1&2 68K set in algorithmically defined classes of *BRCA1* sequence variants that correlate well with pathogenicity. To provide a point of reference for this statement, Table 1.1 uses the ERS to show that, compared to non-spliceogenic silent substitutions, the B1&2 68K set contained an excess of missense substitutions falling within the key RING and BRCT domains (combined P<2.7×10^-4^), and a strong excess of nonsense substitutions overall (P<1×10^-10^).

**Table 1.1.**
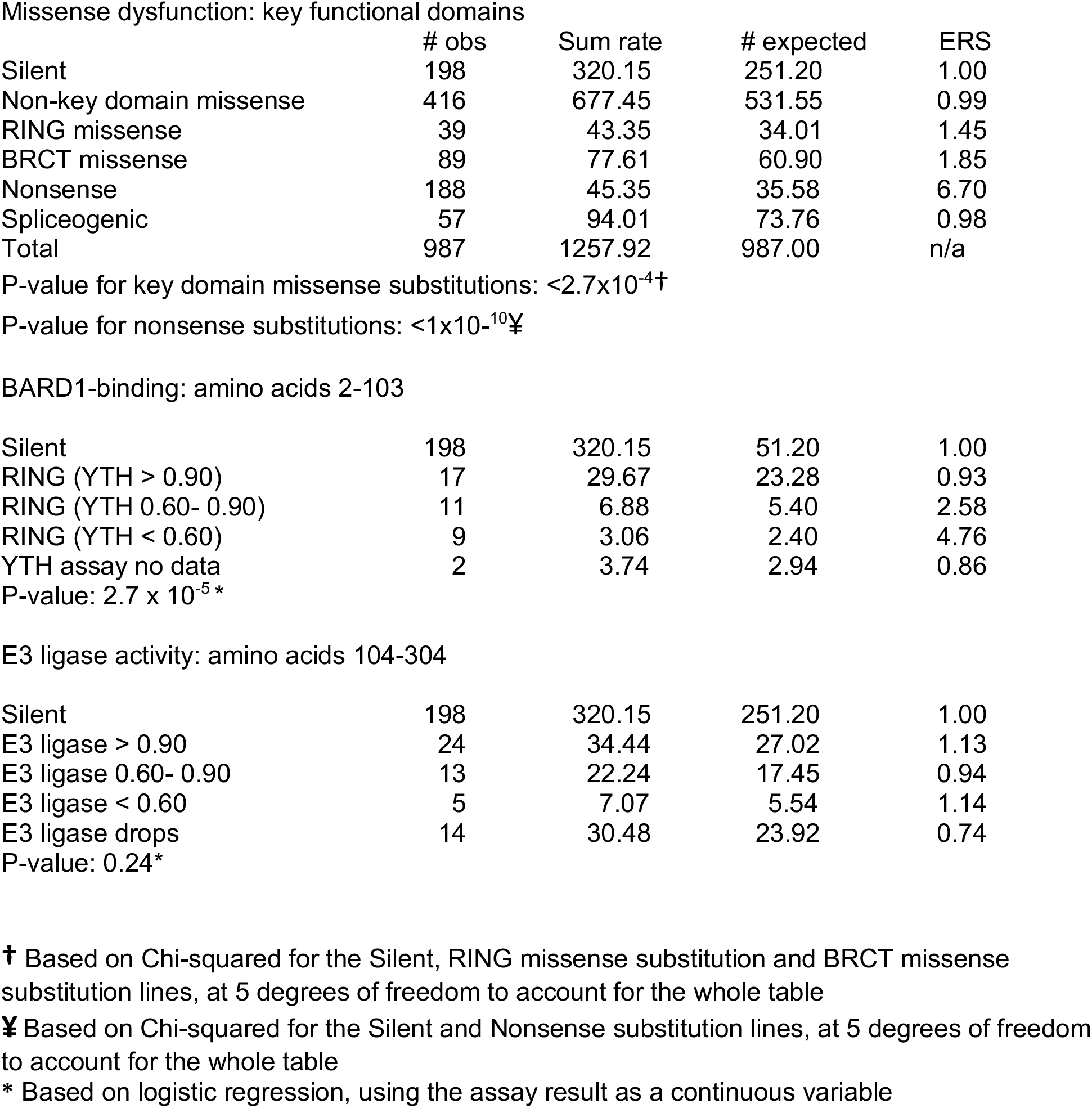
Re-analysis of Starita et al., (2015) yeast 2-hybrid and E3 ubiquitin ligase assays.

Missense dysfunction: key functional domains
|  | # obs | Sum rate | # expected | ERS |
| --- | --- | --- | --- | --- |
| Silent | 198 | 320.15 | 251.20 | 1.00 |
| Non-key domain missense | 416 | 677.45 | 531.55 | 0.99 |
| RING missense | 39 | 43.35 | 34.01 | 1.45 |
| BRCT missense | 89 | 77.61 | 60.90 | 1.85 |
| Nonsense | 188 | 45.35 | 35.58 | 6.70 |
| Spliceogenic | 57 | 94.01 | 73.76 | 0.98 |
| Total | 987 | 1257.92 | 987.00 | n/a |

P-value for key domain missense substitutions: $<2.7 \times 10^{-4}$ †
P-value for nonsense substitutions: $<1 \times 10^{-10}$ ¥

BARD1-binding: amino acids 2-103
|  |  |  |  |  |
| --- | --- | --- | --- | --- |
| Silent | 198 | 320.15 | 51.20 | 1.00 |
| RING (YTH > 0.90) | 17 | 29.67 | 23.28 | 0.93 |
| RING (YTH 0.60- 0.90) | 11 | 6.88 | 5.40 | 2.58 |
| RING (YTH < 0.60) | 9 | 3.06 | 2.40 | 4.76 |
| YTH assay no data | 2 | 3.74 | 2.94 | 0.86 |
| P-value: $2.7 \times 10^{-5}$* | | | | |
| E3 ligase activity: amino acids 104-304 |  |  |  |  |
| Silent | 198 | 320.15 | 251.20 | 1.00 |
| E3 ligase > 0.90 | 24 | 34.44 | 27.02 | 1.13 |
| E3 ligase 0.60- 0.90 | 13 | 22.24 | 17.45 | 0.94 |
| E3 ligase < 0.60 | 5 | 7.07 | 5.54 | 1.14 |
| E3 ligase drops | 14 | 30.48 | 23.92 | 0.74 |
| P-value: 0.24* |  |  |  |  |

† Based on Chi-squared for the Silent, RING missense substitution and BRCT missense substitution lines, at 5 degrees of freedom to account for the whole table
¥ Based on Chi-squared for the Silent and Nonsense substitution lines, at 5 degrees of freedom to account for the whole table
\* Based on logistic regression, using the assay result as a continuous variable

Framing a hypothesis test of the Starita et al. (2015) Y2H functional assay as a predictor of pathogenicity, when these data from Asp2-Asn103 were stratified into substitutions with relatively high (>90% of wt), moderate (60%-90% of wt), and low (<60% of wt) *BARD1* binding activity, the moderate category had an elevated ERS and the low category had a markedly elevated ERS, approaching that of nonsense substitutions (Table 1.1). Using the Y2H % wt activity as a continuous variable, logistic regression confirms that the probability of pathogenicity increases as the Y2H interaction decreases, with P=2.7×10^-5^ (Table 1.1).

Framing a hypothesis test of the Starita et al. (2015) E3 ubiquitin ligase assay as a predictor of pathogenicity for missense substitutions *outside* of the BARD1 interaction domain of *BRCA1*, when these data from Ser104-Glu300 were stratified into substitutions with relatively high (>90% of wt), moderate (60%-90% of wt), and low (<60% of wt) E3 ubiquitin ligase assay, neither the moderate nor the low category had a notably elevated ERS (Table 1). Using these E3 ligase activity data as a continuous variable, there was no relationship between E3 ligase activity and probability of pathogenicity, P=0.24 (Table 1.1).

A secondary question raised by Starita et al is whether the E3 ubiquitin ligase assay adds predictive power over and above the Y2H assay for missense substitutions within the BARD1 interaction domain. This was tested by adding the E3 ubiquitin ligase assay results from Asp2-Asn103 to the Y2H logistic regression. The analysis indicates that the probability of pathogenicity increases as both the Y2H interaction decrease and E3 ubiquitin ligase activity decrease, with P=1.1×10^-2^ for Y2H and 6.7×10^-3^ for E3 ubiquitin ligase activity.

### Mammalian 2-Hybrid Results

The *BRCA1* Ex-UV database (http://hci-exlovd.hci.utah.edu/home.php?select_db=BRCA1) records seven Definitely Pathogenic and one Likely Pathogenic missense substitution in the RING domain. These eight substitutions all had <5% wt activity in the mammalian 2-hybrid BARD1 interaction assay (M2H assay) (Fig. 1.1A). The database records two Not Pathogenic missense substitutions, and these both had >55% wt activity in the M2H assay (Fig. 1.1A, with underlying data in Supp. Table S1.1).

**Figure 1.1.**
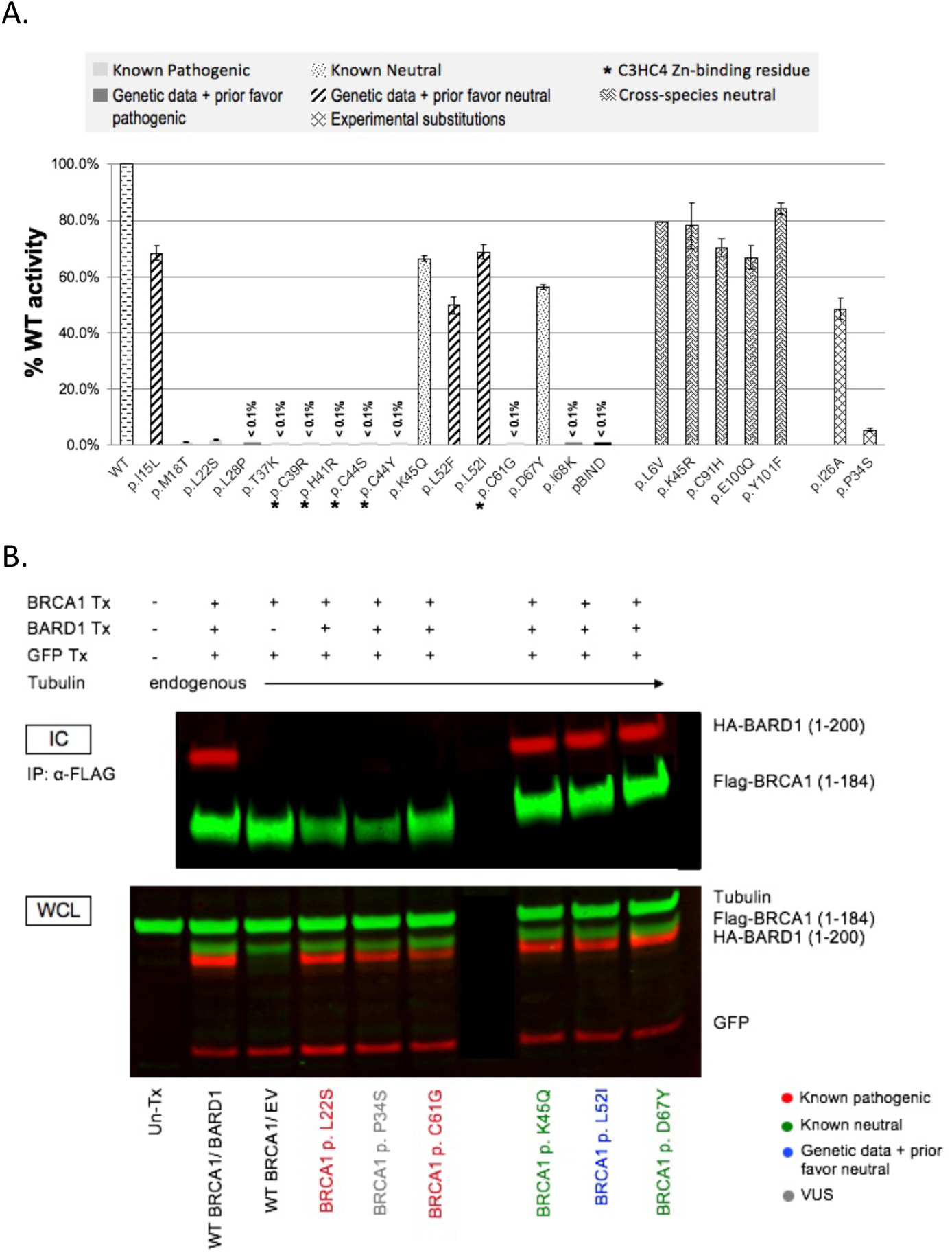
Mammalian two-hybrid BRCA1:BARD1 heterodimerization results. The classification of each substitution assayed is given above the bar graph. Error bars represent 95% confidence intervals. Note that the cross-species neutral variant (p.L6V) does not have a confidence interval because only two (instead of 3 or more) valid expression constructs were made from this variant. **B**. BRCA1:BARD1 co-immunoprecipitation results. Inclusion of BRCA1, BARD1, and/or GFP in the transfections is indicated above the Western blot panels. The identity of the BRCA1 expression construct included in the transfection is indicated below the panels. The upper panel is an immunoblot of immune precipitated immunocomplexes (IC). The lower panel is an immnunoblot of whole cell lysates (WCL). The identity of each protein band is indicated at the right side of the panels.

Although the M2H assay produced clear separation between known pathogenic and known neutral missense substitutions, the total number of known classified substitutions is too few, and too skewed towards pathogenic substitutions, to generate a reasonable calibration curve. Accordingly we assayed 10 more substitutions, systematically selected from three groups as follows. First, the B1&2 68K set included one additional substitution (p.I68K) with observational odds >2:1 in favor of pathogenicity. Second, that data set included four additional substitutions (p.I15L, p.L28P, p.L52F, and p.L52I) with observational odds <0.5:1 in favor of pathogenicity. Third, because substitutions with data in favor of pathogenicity outnumbered those with data against pathogenicity, we identified from the BRCA1 protein multiple sequence alignment used for generating Align-GVGD scores (Tavtigian et al. 2008) (http://agvgd.hci.utah.edu/BRCA1_Spur.html) cross-species amino acid substitutions that met a double criterion indicative of neutrality: (i) the alternate amino acid was present in a primate sequence, and (ii) even if the primate sequence with one of these substitutions is removed from the alignment, the alternate amino acid remains within the range of variation of the mammals-only alignment. Five substitutions met these criteria: p.L6V, p.K45R, p.C91H, p.E100Q, and p.Y101F. The M2H activities of these 10 substitutions are also displayed in Figure 1.1A and, for nine of these ten, the patient derived Odds_Path, missense analysis Prior_P, and M2H result were all congruent. One, p.L28P, presented a more complex pattern: the observational Odds_Path were 0.42 (more than two-fold against pathogenicity), the missense analysis Prior_P was 0.81 (strongly in favor of pathogenicity), and the M2H results was 0.08% of wt activity. Since a Bayesian combination of the Prior_P and Odds_Path result in a Post_P of 0.64 (above 0.5, therefore in favor of pathogenicity), we also interpret the M2H result as congruent.

Therefore, across 20 missense substitutions, the sensitivity and specificity of this M2H assay were both 100% (for both, the 95% CI was 0.69-1.00).

For a subset of variants, the 2-hybrid data were validated at the protein level. The N-terminus of *BRCA1* (amino acids 1-184), fused to FLAG, was immune purified from whole cell lysates. Co-immunoprecipitation of BARD1 (amino acids 1-200), fused to hemagglutinin (HA), was determined by immunoblot. In three separate pull downs, wt, two known neutral (*BRCA1* p.K45Q and p.D67Y), and a *BRCA1* RING missense substitution with genetic data and missense analysis Prior_P against pathogenicity (*BRCA1* p.L52F) bind to BARD1 (Fig. 1.1B). In contrast, the empty vector control and two pathogenic RING missense substitutions (*BRCA1* p.L22S and p.C61G) show no detectable level of BARD1 in their pull downs (Fig. 1.1B). In the assays where no interaction was detected, the presence of BARD1 in the corresponding whole cell lysates confirms that the absence of an interaction in the immune complexes is not due to protein degradation.

### Calibration of the mammalian 2-hybrid assay

A simple regression was used to convert % wt activity from the M2H assay to Odds_Path (Functional LR), the variable required to perform a Bayesian integration of the functional assay data with patient-derived Odds_Path and/or the missense analysis Prior_P (Vallée et al. 2012). For the 15 substitutions with patient observational data, we regressed the % wt activity against log_10_(Odds_Path), where the latter data were obtained either from the Ex-UV database or else from the B1&2 68K set data set. For the 5 cross-species neutral variants, we replaced patient observational data with the missense analysis Prior_P (0.03 <http://priors.hci.utah.edu/PRIORS/> and underlying algorithms), converted to log_10_(Odds).

With % wt M2H activity expressed as a decimal, the calibration equation resulting from the regression is:

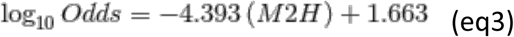

with a 95% confidence interval on the slope of (-6.02 - -2.76) and P=1.78×10^-5^ against the null hypothesis of no relationship between M2H activity and patient observational Odds_Path. The estimated tipping point – the point at which M2H activity switches from evidence against pathogenicity to evidence in favor of pathogenicity, comes a 37.85% of wt activity. From the regression curve and M2H data displayed in Figure 1.2, the interpretive difficulty is that there were no M2H data between 5% and 45% of wt activity. Consequently, the shape of the regression curve through the tipping point is imposed by the (logistic) regression chosen. A simple approach to building this uncertainty into operational conversion from M2H activity to a Functional LR is to use the width of the 80% confidence envelope (80% CE) of the regression to constrain the Functional LR, introducing a “grey zone” of no information around the tipping point, and the upper and lower 80% confidence envelopes to moderate the regression conversion above and below.

**Figure 1.2.**
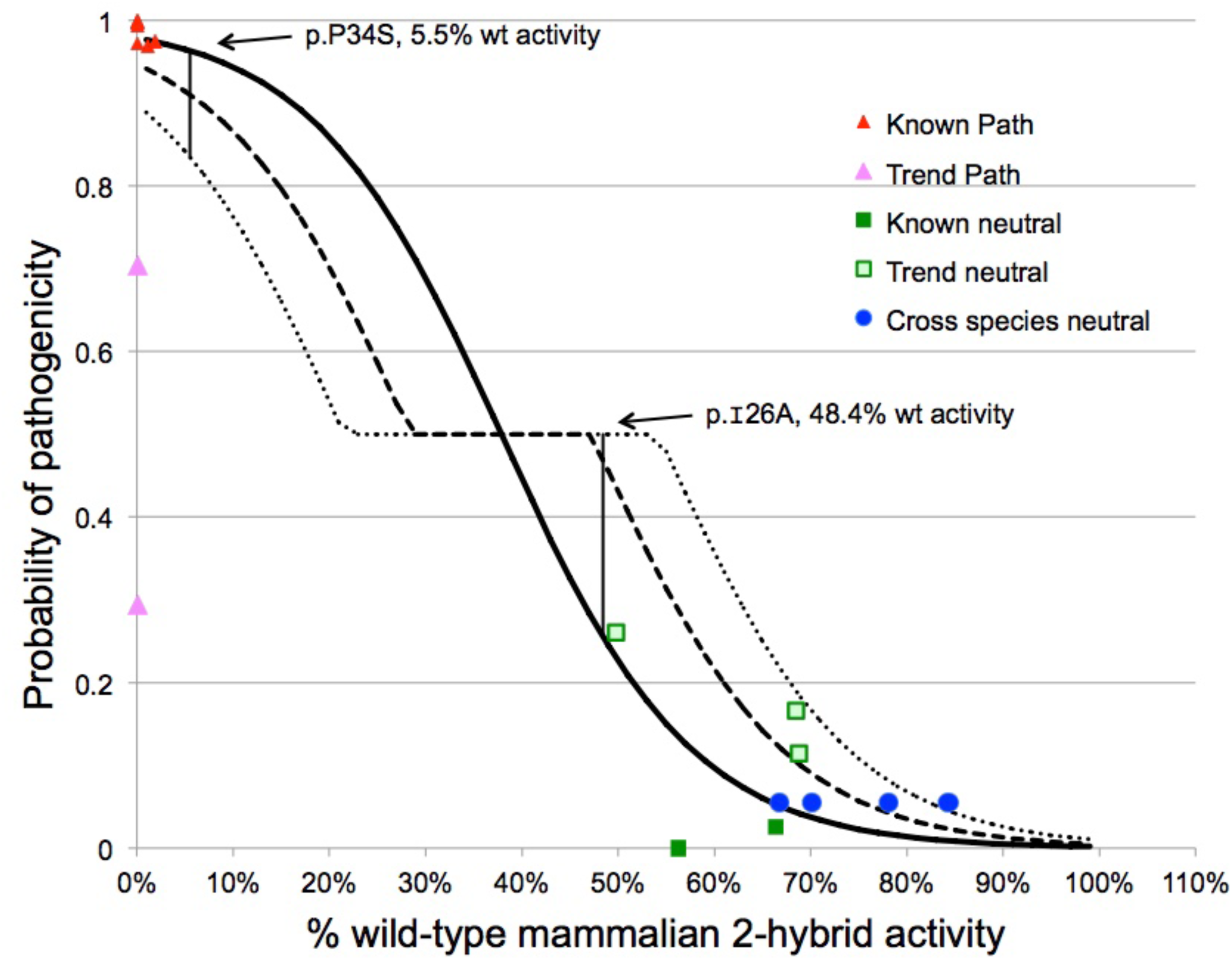
Calibration of the mammalian 2-hybrid assay. Solid line represents the calibration equation resulting from regression of % wt activity against log10(Odds_Path), with the Y-axis expressed as probability instead of odds. The heavy dashed line represents the regression equation modified using the 80% confidence interval of the slope where the regression crosses probability=0.50. The lighter dashed line represents the regression equation modified using the 95% confidence interval of the slope where the regression crosses probability=0.50. The classes of variants included in the regression are indicated in the legend at the right of the figure. The % wt activity of p.P34S (5.5%) and p.I26A (48.4%) are annotated on the figure.

From the regression, we can estimate several additional variables including the mean standard error (MSE), the standard error of each value to be predicted, (*s*^2^{ *pred*}), and W, obtained from W^2^=2F, where F is the upper 80% quantile from the F-distribution with 2 and n-2 degrees of freedom. From the regression MSE=1.333 and W=1.882. Using these to calculate Working-Hotelling 80% confidence band limits (Working and Hotelling, 1929), the moderated regression conversion to the Functional LR becomes: (eq4)

if M2H< 0.22, lower 80% CE, 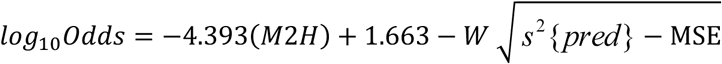

if 0.27 ≤ M2H ≤ 0.50, grey zone, *log_10_ Odds* = *0.00*

if M2H> 0.54, upper 80% CE, 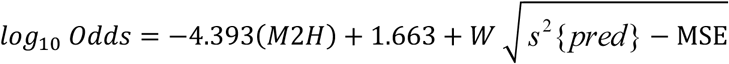

With conversion for *log_10_Odds* to probability of pathogenicity, these are displayed in Figure 1.2.

### Experimental missense substitutions

Two of the missense substitutions evaluated with the M2H assay, p.I26A and p.P34S, were considered experimental rather than part of the calibration series. p.I26A, which disrupts BRCA1:BARD1 E3 ligase activity but does not notably increase cancer susceptibility when homozygosed in mice (Brzovic et al. 2003; Shakya et al. 2011), had 48.4% of wt M2H activity. This is at the lower bound of the known neutral substitutions, but far above the activity of the known pathogenic substitutions.

*BRCA1* p.P34S was observed in a woman diagnosed with ovarian cancer at age 55 and shared by her older sister who was diagnosed with ovarian cancer at age 75 (Figure 1.3). The M2H assay revealed that the substitution had 5.5% of wt activity; this is at the upper bound of the known pathogenic substitutions, but far below the activity of the known neutral substitutions. Quantitative Integrated Evaluation combining the missense analysis Prior_P, segregation, co-occurrence, and M2H constrained Functional LR resulted in either IARC Class 5 or Class 4 (Pathogenic or Likely Pathogenic), depending on the M2H calibration used (Table 1.2).

**Figure 1.3.**
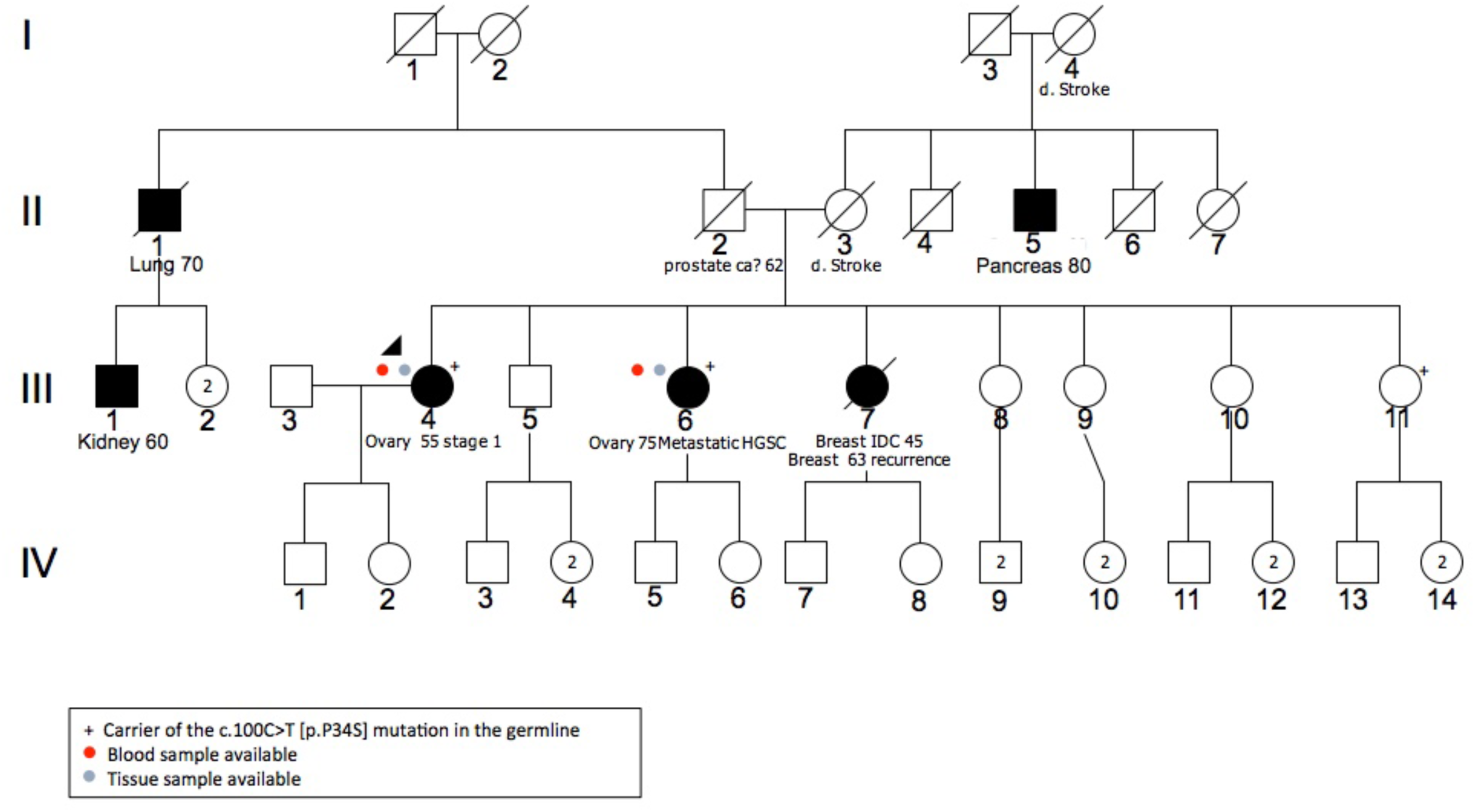
Pedigree with BRCA1 p.P34S. The index case is III-4, diagnosed with ovarian cancer at age 55 years. “+” indicates mutation carriers. Cancer diagnosis and age of onset are indicated for affected family members.

**Table 1.2.** Quantitative Integrated Evaluation of *BRCA1* p.P34S

|  |  |  |
| --- | --- | --- |
| Gene | BRCA1 |  |
| Accession# | NM_007294.3 |  |
| Exon | 3 |  |
| HGVS DNA change | c.100C>T |  |
| BIC DNA change | 219C>T |  |
| HGVS protein change | p.P34S |  |
| Missense Prior_P† | 0.81 |  |
| Splicing Prior_P | 0.02 |  |
| Combined Prior_P | 0.81 |  |
| Segregation LR | 1.77 |  |
| Pathology LR | - |  |
| Sum Family LR | - |  |
| Co-occurrence LR | 1.03 |  |
|  | Functional LR from: |  |
|  | Regression | 80% CE |
| Functional Assay LR* | 26.3 | 6.15 |
| Product of LRs | 47.9 | 11.2 |
| Post_P | 0.995 | 0.980 |
| IARC Class | 5 | 4 |

<sup>†</sup> This is a non-conservative substitution falling at a key domaininvariant position in the reference BRCA1 protein multiple sequence alignment. The Align-GVGD Class is C65, resulting in the missense substitution severity Prior\_P of 0.81.
\* P.P34S had 5.5% of wild-type activity in the calibrated mammalian 2-hybrid assay.

### Towards a high-throughput M2H assay

In principle, recently developed technologies could be used to convert the 1-by-1 M2H assay to a massively parallel high-throughput assay. One strategy would chain together array synthesis and *en masse* Gibson assembly to generate libraries with massive numbers of systematically designed sequence variants; Flp-In to convert *en masse* library transfection to cells with a unique expression construct; two or more colors of 2-hybrid reporters (at different loci) to increase resolution across the range of BRCA1:BARD1 heterodimerization activity; and massively parallel sequencing across bins of multi-color flow sorted cells to read out the M2H activity of individual missense substitutions (Fig. 1.4).

**Figure 1.4.**
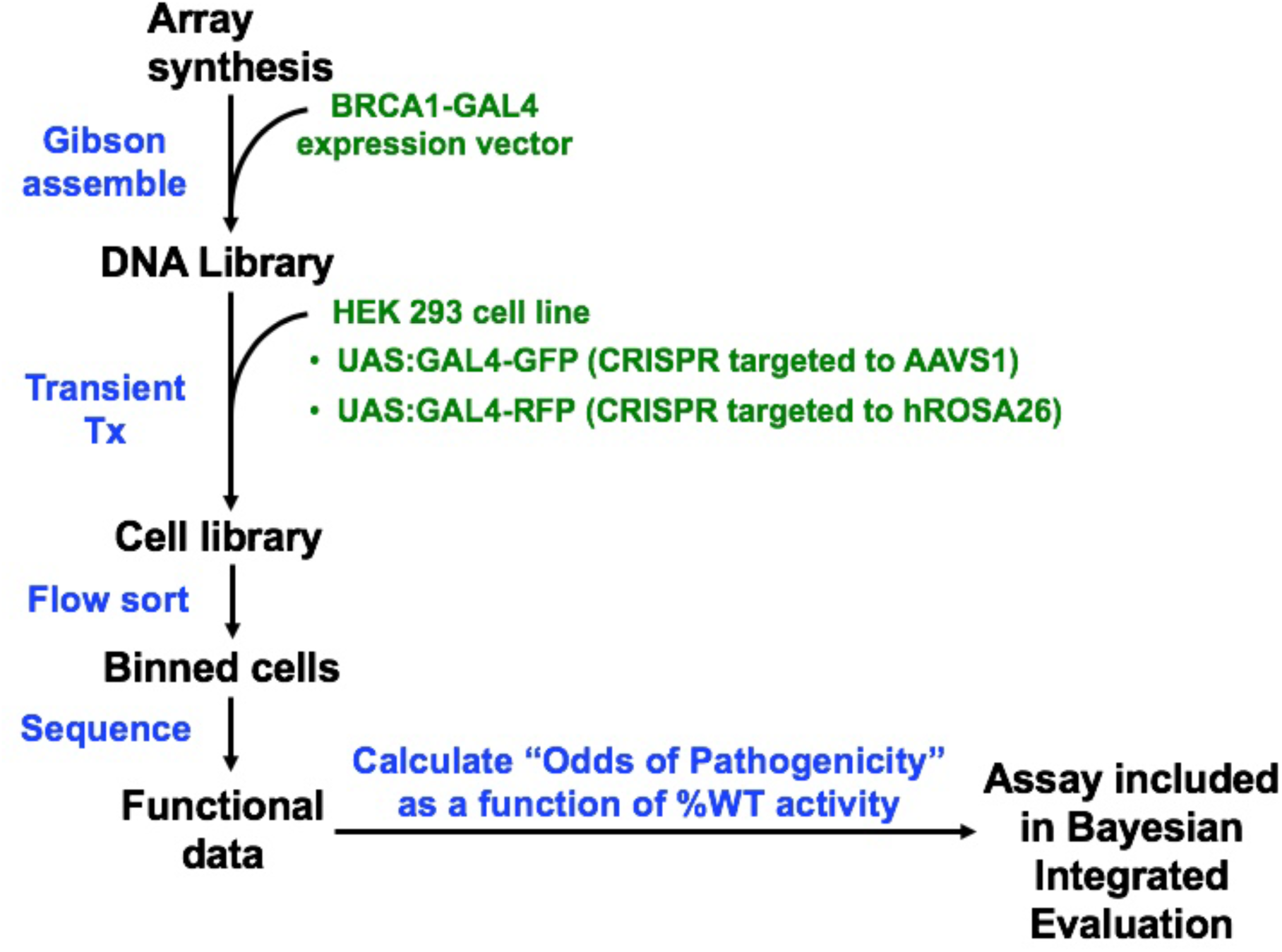
Flow diagram for a potential high-throughput BRCA1:BARD1 RING domain mammalian2-hybrid assay.

To simulate the 2-color reporter, flow sorting, and sequencing phases of this strategy, we prepared a pool of HEK293 cells with UAS GAL4-driven ZsGreen (GFP) and Tdtomato (RFP) M2H reporters. A preliminary test showed that about 12% of the cells in the pool could productively express the red reporter, and about 8% could productively express the green reporter (data not shown). Individual wells of a 6-well plate were then transiently co-transfected with wt, two neutral, two pathogenic and one VUS (now re-classified as Likely pathogenic) *BRCA1* RING missense substitution (plus wt *BARD1*). Post transfection, the six wells of cells were mixed and then FACS sorted into four bins based on RFP expression. Each of these bins was further sorted into four bins based on GFP expression.

Sequencing of *BRCA1* RING domain transgene cDNA from each of the four bins along the dual-expression main diagonal (Red_low_:Green_low_ => Red_high_:Green_high_) (R_L_G_L_ => R_H_G_H_), revealed that 80% of *BRCA1* reads aligned perfectly to one of the six possible target sequences. Table 1.3 summarizes read counts and analyses, restricted to the perfect-match reads, of these data. We observed considerable separation in % wt activity between the two neutral and two pathogenic substitutions included in the experiment. Moreover, the experiment exactly recapitulated the ordering of % wt activity that we obtained in the 1-by-1 M2H assay. Finally, placing the two pathogenic rare missense substitutions {p.L22S+ p.C61G} in one category and the two neutral substitutions {p.K45Q+ p.D67Y} in a second category, a simple 2-sided T-test revealed P=0.029 against the null hypothesis that the two categories had equal representation in the R_H_G_H_ versus R_L_G_L_ bins.

**Table 1.3.** Mammalian 2-hybrid dual color flow sort and sequencing assay

| Missense Substitution | 1-by-1 M2H %wt activity | FACS and sequencing: perfect match read counts¥ |  |  |  | FACS & seq % wt activity† |
| --- | --- | --- | --- | --- | --- | --- |
|  |  | Bin A1 | Bin B2 | Bin C3 | Bin D4 |  |
| Wild-type | 100.0 (ref) | 418,971 | 517,025 | 585,459 | 347,951 | 100.0 (ref) |
| p.K45Q | 66.4 | 298,650 | 413,076 | 339,096 | 259,568 | 104.5 |
| p.D67Y | 56.3 | 221,276 | 294,912 | 277,169 | 156,203 | 85.5 |
| p.P34S | 5.51 | 178,460 | 112,849 | 61,752 | 21,700 | 15.1 |
| p.L22S | 1.92 | 124,040 | 108,186 | 50,912 | 11,375 | 11.4 |
| p.C61G | 0.03 | 101,911 | 19,560 | 2,567 | 604 | 0.7 |

¥ A, B, C, and D indicate increasing red channel signal and 1,2,3, and 4 indicate increasing green channel signal. Thus A1, B2, C3, and D4 are the main diagonal for increasing signal in both channels.
† estimated from read counts as $[(\text{BinD4})/(\text{BinA1}+\text{BinD4})_{\text{experimental}}]/[(\text{BinD4})(\text{BinA1}+\text{BinD4})_{\text{wildtype}}]$ .

## DISCUSSION

Structural and biochemical studies of the BRCA1 E3 ubiquitin ligase activity, including recent evidence that BARD1 residue Arg99 is critical to this activity, put BRCA1:BARD1 heterodimerization upstream of E3 ubiquitin ligase activity towards key substrates (Brzovic et al. 2003; Densham et al. 2016). This heterodimerization is also required for early recruitment of *BRCA1* to sites of DNA double strand breaks (Li and Yu, 2013). While it is not clear how the various activities attributed to BRCA1, heterodimerization dependent or not, add up to the protein’s full tumor prevention activity, it is more likely than not that heterodimerization is the key function residing in the C3HC4 RING finger and helical bundles that constitute the first ∼100 amino acids of this protein. In contrast, evidence that the ubiquitin ligase deficient substitution *BRCA1* p.I26A does not cause notable tumor susceptibility in mice, combined with evidence against pathogenicity in humans for ubiquitin ligase damaging substitutions falling immediately downstream of the RING domain, weigh against the hypothesis that ubiquitin ligase activity is central to the full tumor prevention activity BRCA1.

The high-throughput Y2H assay described by Starita et al. (2015) and the 1-by-1 mammalian 2-hybrid assay are both very nearly direct tests of BRCA1:BARD1 heterodimerization activity. While both assays are able to resolve the activity of substitutions falling at the key C3HC4 RING cysteines and histidine from neutral substitutions, there was a notable difference in their discrimination between other pathogenic substitutions and neutral substitutions. From the data underlying Figure 1.1, the three known pathogenic substitutions plus two additional substitutions with combined Prior_P and Odds_Path leaning towards pathogenicity that were used for assay calibration averaged 0.66% of wt (SD = 0.84%) while the two neutral substitutions plus three additional substitutions with combined Prior_P and Odds_Path leaning towards neutrality averaged 61.9% of wt (SD = 8.5%). Using the sum of the standard deviations of the pathogenic and neutral substitutions as a yardstick, these two groups were resolved by 6 summed standard deviations. In contrast, from Starita et al. (2015), these two groups averaged 89.1% (SD = 7.2%) and 100.6% (SD = 8.2%) of wt activity, respectively. Thus they were resolved by less than one summed standard deviation, which is indicative of limited assay sensitivity. Rescuing sensitivity by adding the high-throughput E3 ubiquitin ligase assay would introduce a systematic source of error because there genuinely are some missense substitutions that are proficient for heterodimerization but deficient for E3 ligase activity, and there is evidence against pathogenicity for this class of substitutions.

The magnitude of risk conferred by heterozygous pathogenic missense substitutions in *BRCA1*, whether measured as penetrance or odds ratio, is clearly a continuous variable. However, virtually all of the *BRCA1* missense substitutions so far placed in IARC Classes 4 or 5 are thought to be high-risk, essentially equivalent to protein truncating variants (Goldgar et al. 2004) or conferring odds ratios of 5 or higher (Easton et al. 2015). Very few have been classified as moderate-risk; the BRCT substitution *BRCA1* p.R1699Q is the only established moderate-risk *BRCA1* substitution (Spurdle et al. 2012), and no RING domain substitutions have been established as such. If a functional assay were very accurate, there would in principle be a threshold in % wt activity marking the tipping point between evidence for or against pathogenicity, a range corresponding to moderate-risk, and a threshold below which the substitutions are most likely high-risk. However, absent moderate-risk substitutions to include in a calibration, the % wt activity levels corresponding to these thresholds are necessarily uncertain. Here, we used the width of the 80% confidence envelope of the regression equation to introduce a % wt activity “grey zone” at Functional LR = 1.0. We suggest, subject to community discussion, that an 80% CE is appropriate to constrain functional assays thought to be directly related to the underlying mechanism of pathogenicity, and the 95% CE when the assay appears reasonably accurate but the underlying mechanistic connection is less clear. While this approach to constraining the Functional LR to cautiously enable higher throughput VUS classification is somewhat *ad hoc*, the appropriate comparators are the level of *ad hocness* in the way that the ACMG guidelines incorporates functional assays (Richards et al. 2015), and the complexity of the functional assay rules in the InSiGHT mismatch repair gene variant classification criteria (Thompson et al. 2013).

For humans, the *de novo* nucleotide substitution rate is thought to be ∼1×10^-8^ substitutions per site per generation (Shendure and Akey, 2015; Veltman and Brunner, 2012). Allowing that there are 30-fold differences in substitution rates across sites (Lunter and Hein, 2004), this average rate may over-estimate the median rate by 10-fold. Even so, a rate of 1×10^-9^ substitutions per site per generation taken against the human population of 7×10^9^ individuals means that the human gene pool includes multiple substitutions at almost every nucleotide, in turn implying multiple rare missense substitutions at almost every codon, many of which are pedigree-specific because they occurred within the last few generations. For this reason, using a 1-by-1 functional assay to chase after clinically observed substitutions as they are reported only makes sense if there are technical reasons to believe that a high-throughput assay (i) meeting community standards for expression construct assembly and verification, and (ii) possessing sufficient dynamic range to contribute to variant classification in combination with multiple independent lines of evidence, is technically impossible. The expression construct criterion, primarily that each construct be independently prepared three or more times and sequenced (Guidugli et al., 2013; Iversen et al., 2011), can be met by strategies that combine array synthesis, barcoding, and massively parallel sequencing. For missense substitutions in *BRCA1*, the vast majority that have been classified as IARC Class 4 or 5 had missense analysis Prior_Ps of either 0.66 or 0.81. Back calculating from Bayes’ rule, Functional LRs of 9.8 and 4.5, respectively, would suffice to convert those Prior_Ps to Post_Ps of 0.95, just reaching the Class 4 “Likely Pathogenic” criterion (Plon et al. 2008). These values are within the dynamic range of the 1-by-1 M2H assay and may well fall within the dynamic range of a high-throughput version of the same assay.

**Supplemental Table 1.1.**
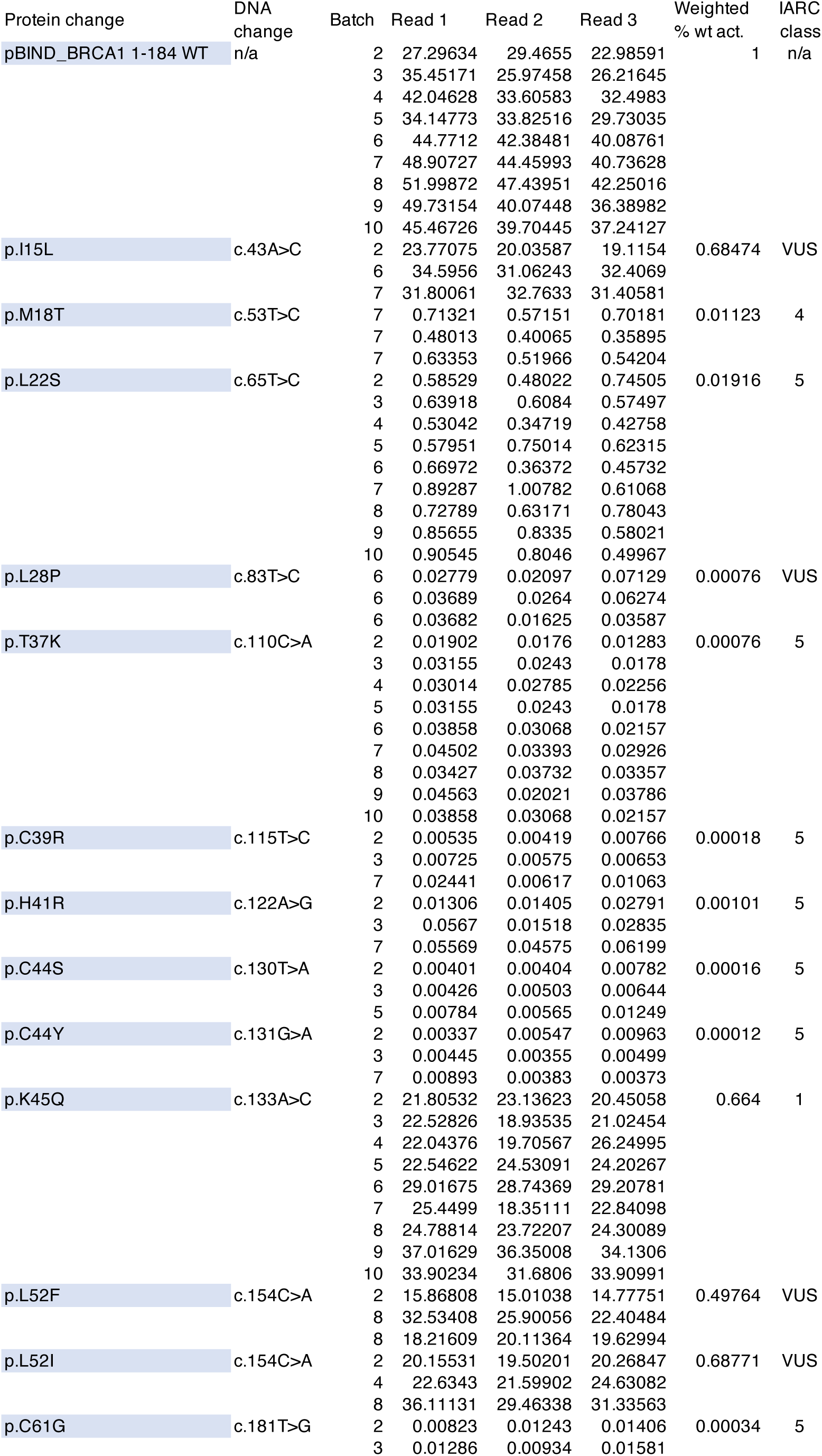
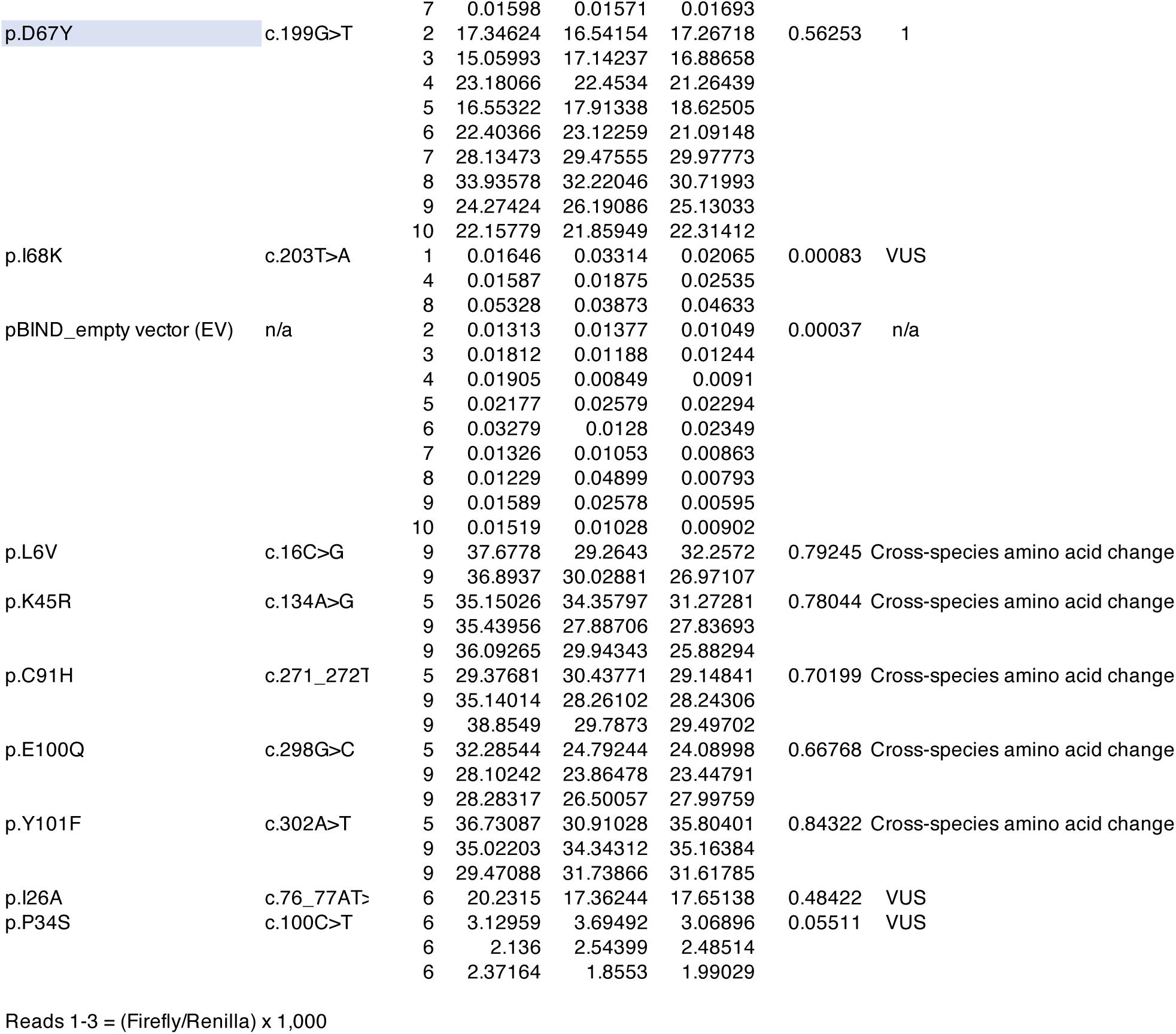

## INTRODUCTION to PART 2

When individuals undergo clinical genetic disease predisposition testing by re-sequencing of high-risk and/or moderate-risk susceptibility genes, most outcomes fall into one of three categories: a pathogenic sequence variant(s) is found, no reportable sequence variant(s) is found, or a variant(s) of uncertain significance (VUS) is found.

Identification of pathogenic variants can modify the management of affected patients and inform selection of risk-reduction strategies for at-risk carriers. On the other hand, when no reportable variants are found, patients and their families are managed based on their relevant personal and family disease histories. However, a variety of clinical problems arise when a VUS (but no pathogenic variant) is found. For example, for carriers of the VUS, the inherent informational uncertainty can raise anxiety. Physicians will sometimes over-interpret VUS –most of which are actually benign – which may lead to over-treatment of healthy individuals who carry the variant. Alternatively, for the subset of VUS that are actually pathogenic, absence of that information denies patients and their at-risk relatives the medical management benefits that can flow from a clear genetic diagnosis.

*BRCA1* is a high-risk susceptibility gene for triple negative breast cancer and for ovarian and peritoneal cancers arising from the Fallopian tubes. The 1,863 amino acid protein encoded by *BRCA1* heterodimerizes with the 777 amino acid protein encoded by its paralog, *BARD1*. Heterodimerization stabilizes both proteins, and the heterodimer participates in DNA repair related activities at DNA double strand breaks, at stalled DNA replication forks, and at unresolved R-loops (for reviews, see (Venkitaraman, 2019; Tarsounas & Sung, 2020)). Rapid recruitment of BRCA1-BARD1 to sites of DNA double strand breaks favors homologous recombination repair (HRR) over more error-prone mechanisms such as non-homologous end joining (NHEJ) and single -strand annealing (SSA).

Both BRCA1 and BARD1 have an amino-terminal C3HC4 RING finger and a carboxy-terminal pair of BRCT domains. In broad mechanistic overview, the RING fingers are responsible for heterodimerization, which in turn is critical for rapid recruitment of BRCA1-BARD1 to sites of damage. The RING heterodimers also constitute an E3 ubiquitin ligase; this can recruit several different E2 ligases to target an incompletely enumerated set of proteins for degradation. The BRCA1 BRCT domains are a phosphopeptide binding module and participate in a wide network of phosphoprotein interactions. BARD1s BRCT domains bind ADP-ribosylated histones at sites of DNA damage, and this activity is critical for rapid recruitment of the heterodimers to sites of DNA damage.

The vast majority of known pathogenic variants in *BRCA1* are, ultimately, either protein truncating variants or large gene rearrangements. Nonetheless, about 10% of known pathogenic variants are either non-spliceogenic missense substitutions or in-frame insertion or deletion mutations (IFIs, IFDs) (Goldgar et al., 2004; Easton et al., 2007). Both qualitative and quantitative methods have been developed for evaluation and clinical classification of missense VUS (Goldgar et al., 2008; Richards et al., 2015). While both share the feature of integrating multiple types of data towards variant classification, neither has become efficient enough to classify a strong majority of VUS missense substitutions in key domains such as the BRCA1 RING domain. Fundamentally, four classes of data are available for missense substitution evaluation: allele frequency estimates in cases and/ or the population; sequence analysis-based computational tools; patient observational data including family cancer history, segregation in pedigrees, and tumor features; and functional assays. Alone or combined, allele frequencies and computational tools are not powerful enough to classify substitutions as Likely Pathogenic (LP) or Pathogenic (P), though they are sufficient to classify substitutions as Likely Benign (LB) or Benign(B) (Richards et al., 2015). Patient observational data are sometimes very powerful, but are not systematically collected. Functional assays may be more powerful than either allele frequency or computational tools, but high-throughput and/or comprehensive assays are not yet well enough calibrated for confident combination with the other data towards clinically applicable variant classification.

Three high-throughput, nearly comprehensive assays that covered the BRCA1 RING domain have been described. Each has its strengths and weaknesses. The phage display BRCA1 auto-ubiquitination assay is very high throughput, but the observation of a separation of function mutation (p.I26A, which has drastically reduced ubiquitination activity, but is not pathogenic) (Shakya et al., 2011; Starita et al., 2015) argues against the utility of the assay. The yeast 2-hybrid assay for BRCA1-BARD1 heterodimerization is both very high throughput and specifically tests the mechanistically key RING domain function (Starita et al., 2015). However, yeast assays have concerning error rates (Carvalho et al., 2009; Lyra et al., 2020), and should be replaced with corresponding Mammalian assays when possible. Finally, a cell viability assay used a high-throughput genome editing approach to test 96.5% of possible single nucleotide substitutions in the *BRCA1* RING and BRCT domains (Findlay et al., 2018). Independent evaluation of this viability assay’s results confirmed high sensitivity and specificity (Lyra et al., 2020); nonetheless, the mechanistic connection between cell viability in this assay and cancer susceptibility in humans is not clear, leaving room for a complementary, mechanistically relevant assay performed in human cells. The goal of this study was to fill this gap with a comprehensive Mammalian two-hybrid assay.

In the original formulation of the American College of Medical Genetics (ACMG) variant classification guidelines (Richards et al., 2015), individual data types such as functional assays (codes PS3 and BS3) were weighted by expert opinion. More recently, we fitted the qualitative ACMG guidelines into a quantitative Bayesian framework and then contributed to development of guidelines for quantitative calibration of functional assays (Tavtigian et al., 2018; Brnich et al., 2019). Even so, non-circular validation of the initial calibration remains challenging because the number of known pathogenic and benign variants is often so small that they are all required to provide accurate fitting during the initial calibration process, and thus cannot be divided into training and test sets. Here, we take advantage of the comprehensive nature of our assay to validate using patient and population observational data that are independent to the calibration variants. The result allows integration with other data via either the ACMG Bayesian framework or the Bayesian points-system that we derived from it (Tavtigian, Harrison, Boucher, & Biesecker, 2020).

## RESULTS

### Assay development and coverage

Previously, we adapted a chemiluminescent Mammalian 2-hybrid (M2H) assay, assessing interaction between the RING domains of BRCA1 and BARD1, for evaluation of BRCA1 RING domain missense substitutions and IFDs. Noting that the assay showed perfect concordance with the then-known pathogenic and benign controls, and also correctly scored the synthetic ubiqitination separation of function variant p.I26A as functional, we adapted the assay to a four-color fluorescent protein format that is compatible with fluorescence activated cell sorting and massively parallel sequencing (Paquette et al., 2018). These adaptations improved the throughput of the approach and enabled more comprehensive assessment of variants in the BRCA1 RING domain.

The RING domain is short enough that RNA-seq can be used to sequence the entire RING coding sequence. We used array synthesis to generate oligos containing all possible missense substitutions from amino acid 2 to 100 that can be created by a single nucleotide substitution. Where two or more nucleotide substitutions create the same missense variant, we generally assayed the substitution predicted to have the highest translation efficiency (n=604). A scan of substitutions to proline (n=95), a scan of frame 0 in-frame single amino acid deletions (n=96), and a scan of in-frame insertions of an alanine residue (n=96) were also assayed (these latter counts are not 99 because there are four prolines, six instances of the same amino acid at consecutive positions, and three alanines in the RING sequence). Oligos containing each sequence variant of interest were array synthesized on a background of silent nucleotide substitutions that assign four distinct “silent barcodes” to each variant. This means that the sequence variants of interest are sequenced, replicates are counted via the number of engineered barcodes observed with each variant, and the only data used for subsequent statistical analyses are sequencing reads with a perfect match to an expected variant-barcode combination, which eliminates the possibility that errors in the sequence confound the interpretation of the results.

The cloning of array-synthesized *BRCA1* fragments, the M2H indicator loci, and transfection are summarized in panels A and B of figure 2.1. In the flow sorting that followed transfection, cells were gated for blue and yellow fluorescence to exclude cells that were not productively transfected with both BRCA1 and BARD1 expression constructs. Activity of the BRCA1 RING sequence variants was assessed by sorting of cells into bins of increasing dual red-green fluorescence followed by RNAseq and read mapping, which is summarized in panel C of figure 2.1. The number of replicates obtained by sequence variant type is summarized in panel A of figure 2.2.

**Fig. 2.1.**
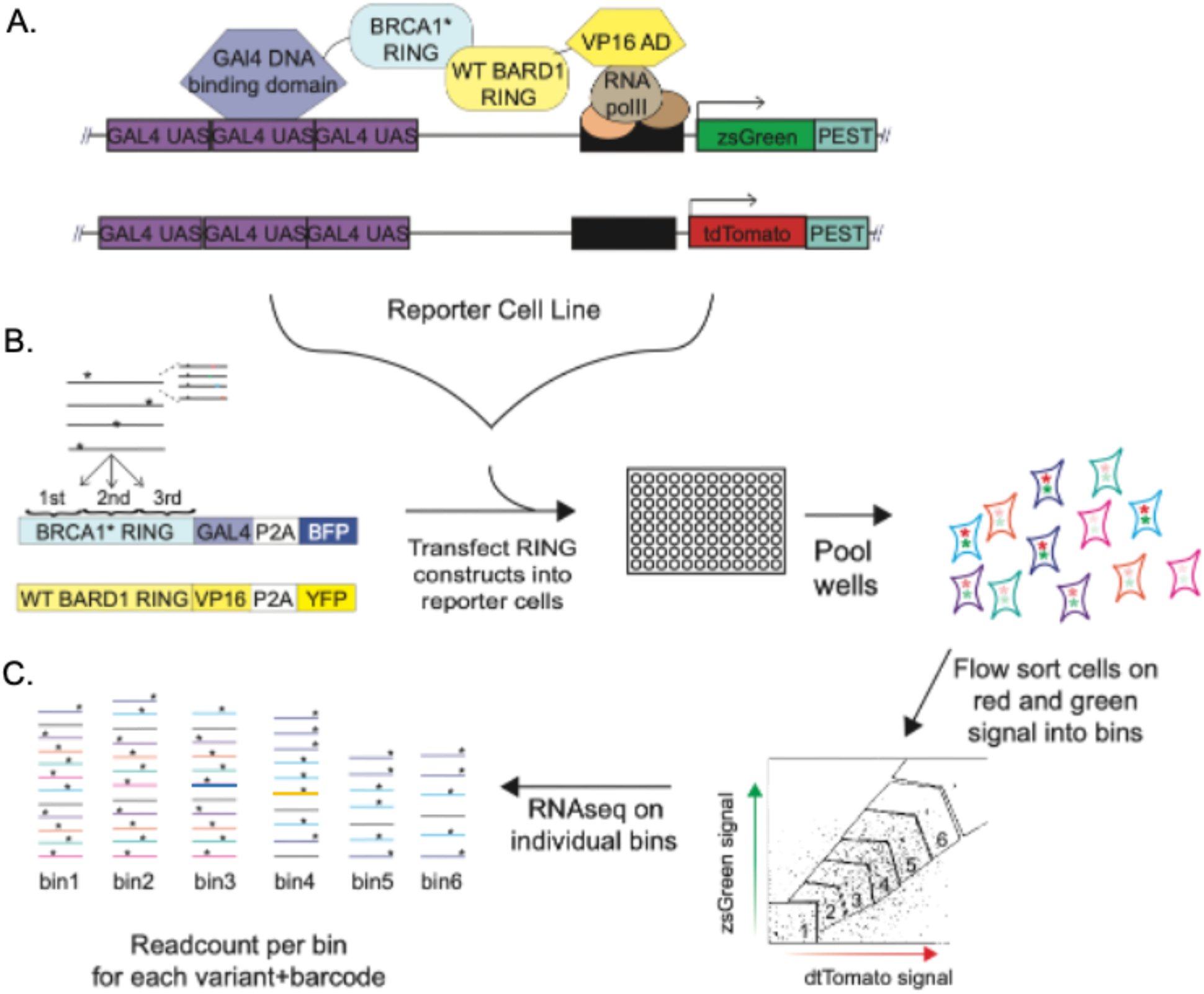
Assay workflow. **A**, Mammalian Two-Hybrid Scheme. The variant-BRCA1-RING GAL4 fusion protein and wildtype-BARD1-RING VP16 fusion protein are shown along with the two integrated GAL-UAS fluorescent reporters. **B**, Workflow for multiplexed assay. 1–Synthesis and cloning mutant oligo array into BRCA1 RING-GAL4-BFP vector backbone. Four versions of each variants are present by use of silent dinucleotide barcodes. The BARD1 RING-VP16-YPF vector is also shown. 2–Transfection of BRCA1 variant-RING-GAL4 constructs into individual wells of reporter cells along with wildtype BARD1 RING-VP16. 3–Transfected wells are trypsinized and pooled. **C**, 1–Cells are flow sorted into 6 bins based on increasing expression of red and green fluorescent reporter activity; 2–RNAseq on BRCA1-RING GAL4 cDNA pools from each bin generates a read count/bin for each variant, reflective of that variant’s activity in the assay. See text for more detailed description of assay.

**Fig. 2.2.**
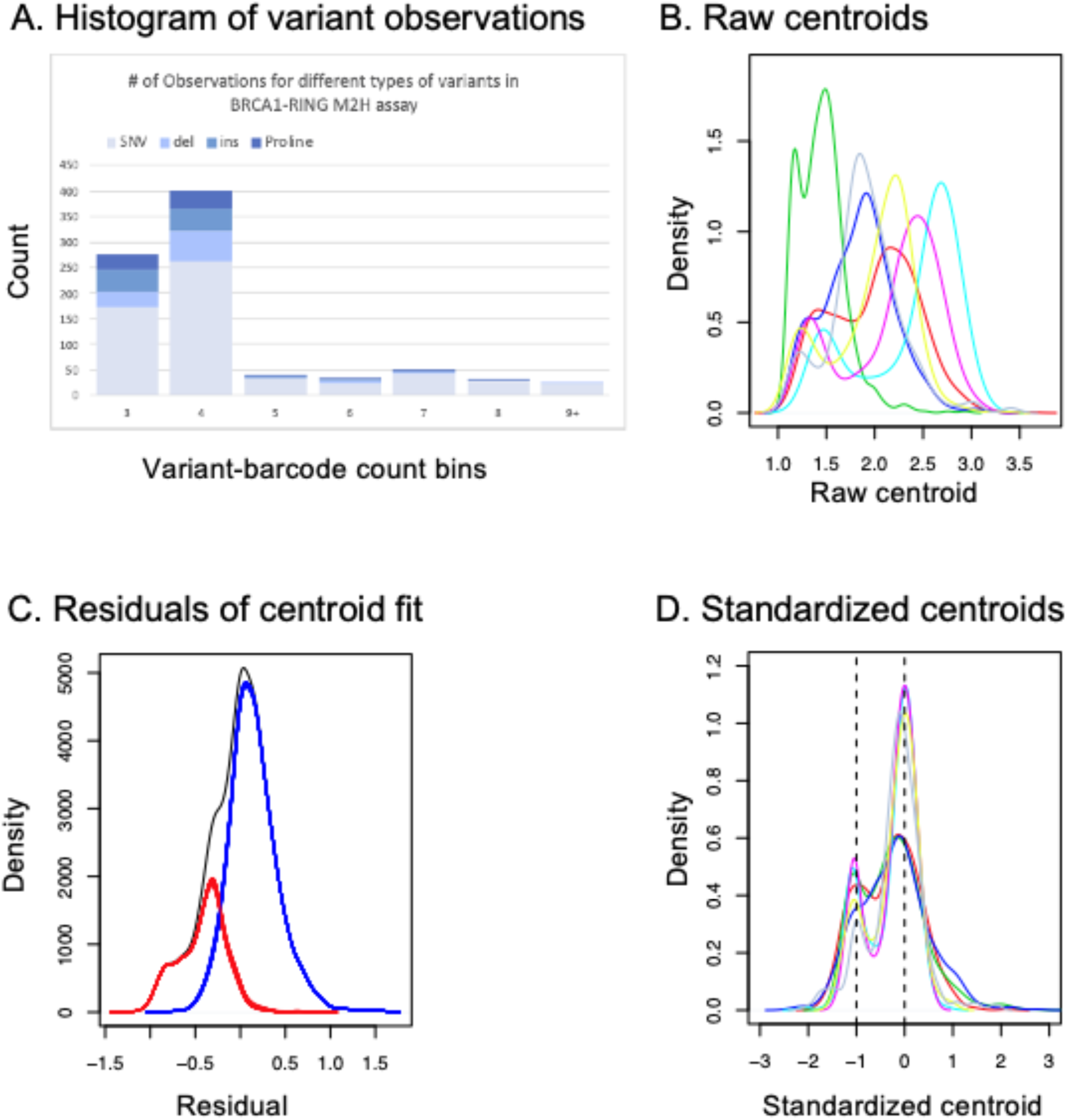
Data merge across the seven experiments. **A**, Histogram of variant barcode observations. **B**, Smoothed centroid density estimates from each of the seven individual experiments. Red: First experiment, first third. Green: First experiment, middle third. Blue: First experiment, final third. Cyan: Second experiment, first third. Magenta: Second experiment, middle third. Yellow: Second experiment, final third. Grey: Final experiment, covering missing or under-represented variants across the entire RING domain. **C**, Distribution of residuals of the centroid fit. Grey: all data. Blue: distribution of residuals attributed to a high activity subset of variants. Red: distribution of residuals attributed to a low activity subset of variants. **D**, Standardized centroids. Color legend as in panel B.

### Data merge across individual experiments

Seven individual experiments were required to drive data acquisition to completion. Panel B of figure 2.2 gives smoothed estimates of the centroid densities for each experiment, showing strong experiment effects and clearly indicating an underlying bimodal Gaussian M2H activity distribution, which we attribute to a dichotomy of the variants into those that have near wild type versus low activity. We first fitted a linear regression for the observed centroid as the dependent variable with factors for experiment, barcode and their interactions, as explanatory variables. The distribution of the residuals for this fit is shown in black in panel C of figure 2.2. This also strongly indicates bimodality. In order to estimate the mixture, mean and variance parameters of a bimodal Gaussian distribution, we then added to the linear regression a factor indicating whether a variant belongs to the upper or lower peak, and all explanatory variable interactions. As this indicator is, of course, not observable, we estimated it and the other parameters using the iterative expectation-maximization (EM) algorithm. For each variant, this yielded both an estimate for the probability of low activity (PLA) and a standardized centroid (SC) that corrects for experiment and barcode effects. The SC distributions for each experiment are shown in panel D of figure 2.2. Using the PLA to randomly assign the variants to the upper or lower peak each time, we then fitted multiple instances of the full linear regression. Density estimates for the residuals from these fits are also shown in panel C of figure 2.2, the red lines giving the residuals for the variants randomly assigned to the lower peak and the blue lines those assigned to the upper. These clearly show the indicator accounting for the bimodality, and that it is robust to the randomization representing the uncertainty regarding peak assignment. At this stage of the analysis, the maximum likelihood estimate for the proportion of variants that fall in the lower peak of the mixture distribution, as obtained from the EM algorithm, was 27.4%.

### Assay calibration

Under the Bayesian interpretation of the ACMG sequence variant classification guidelines, the M2H assay should preferentially be calibrated to provide odds in favor of pathogenicity (Odds_Path) for each variant assayed. To avoid downstream circularities, the calibration should use patient or population observational data (segregation, personal and family cancer history, allele frequency) rather than indirect data that themselves need to be calibrated (e.g., other functional assays, sequence analysis-based computational tools). Starting from a BRCA1/2 key domain prior probability of pathogenicity of 0.35 (Easton et al., 2007; Li et al., 2020), we identified 13 missense substitutions reachable by a single nucleotide substitution (snMS) plus one IFI with patient/population observational evidence sufficient to surpass the LP boundary (Post_P>0.95, Odds_Path>35.3) plus two additional snMS with 0.90<Post_P≤0.95, Odds_Path>16.7 (table 2.1) (Vallee et al., 2012; Easton et al., 2007; Tavtigian, Byrnes, Goldgar, & Thomas, 2008; Spearman et al., 2008; Sweet, Senter, Pilarski, Wei, & Toland, 2010; Whiley et al., 2014; Parsons et al., 2019; Li et al., 2020). Of these, one snMS (c.211A>G, p.R71G) was predicted to damage mRNA splicing and then shown to be pathogenic on that basis (Vallee et al., 2016; Parsons et al., 2019); consequently, p.R71G was excluded from the calibration calculations. We also identified 8 snMS with patient/population evidence sufficient to surpass the LB boundary (Post_P<0.05, Odds_Path<0.0977) plus four additional snMS with 0.10>Post_P≥0.05, Odds_Path<0.206 (table 2.1). Noting a relative excess of variants with evidence of pathogenicity in the calibration set, we identified in the Mammalian subset of our reference *BRCA1* protein multiple sequence alignment (Tavtigian et al., 2008) two evolutionarily tolerated alternate amino acids that remained within the range of variation observed in the alignment even after the first sequence in which they occur is removed: p.K45R and p.C91H. Under the assumption that these are very likely benign, they were added to the calibration set with assigned Odds_Path of 0.206, resulting in n=15 variants with evidence of pathogenicity and n=14 with evidence of benignity.

**Table 2.1.**
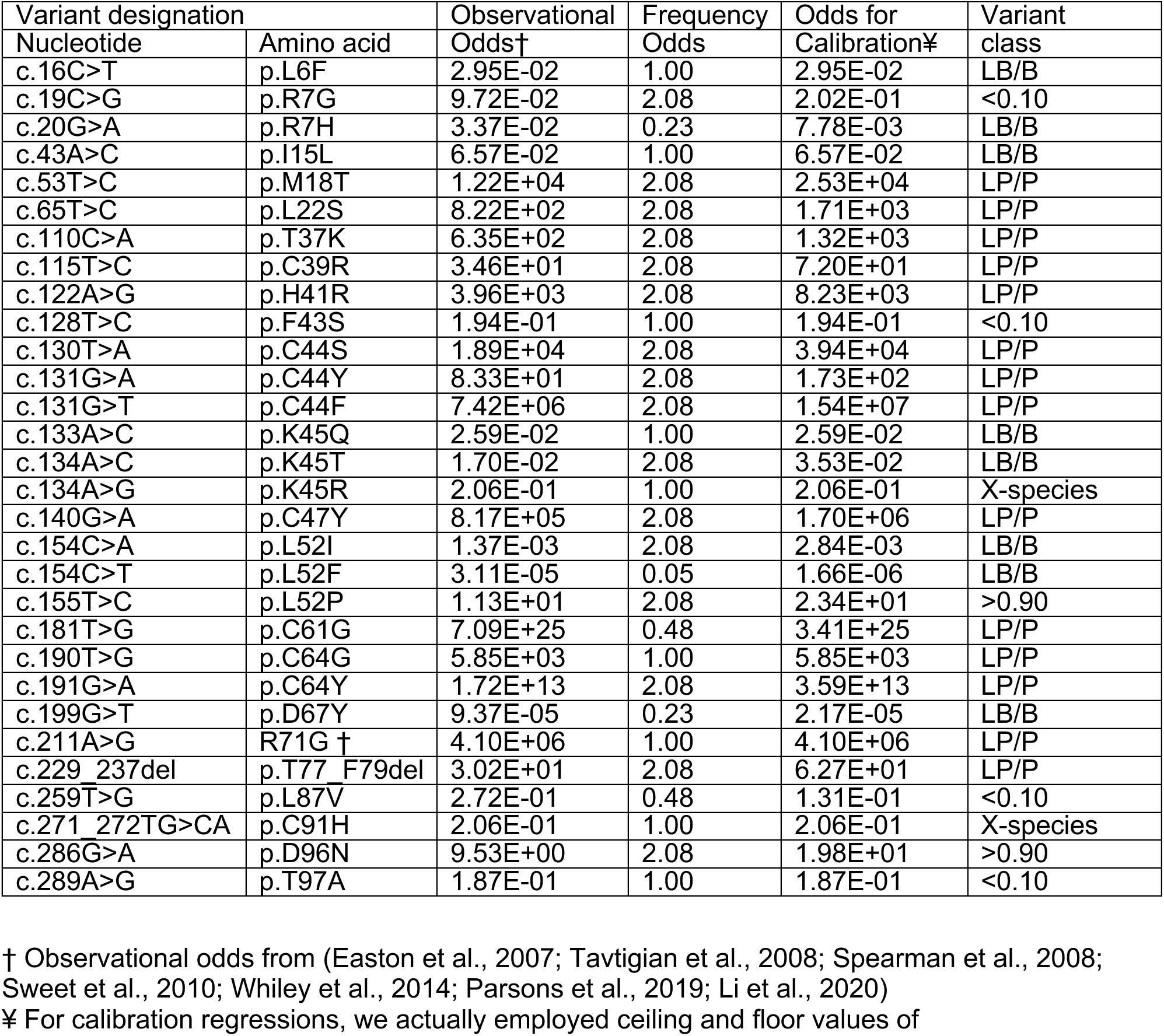
Sequence variants used for Mammalian 2-hybrid assay calibration

Using the probabilities of pathogenicity of the 29 calibrant variants as weights, we estimated the mean vector and variance-covariance matrix of two bivariate Normal distributions for SC and logit(PLA): one bivariate distribution for benign variants and one for pathogenic. The logit, or log-odds, transformation is the standard transformation used to model observations of probabilities as Normals. By applying these estimated probability density functions to the observed SC and logit(PLA) for any variant, we can obtain the variant’s calibrated log likelihood ratio in favor of pathogenicity. In all six panels of figure 2.3, we plot SC versus logit(PLA), and show critical contours of the log likelihood ratio surface. The contours are plotted at values of 81^!/#^ for % = ±1, ±2, and ±4, which are the thresholds of the ACMG Supporting, Moderate, and Strong evidence bins (positive exponents for pathogenic, negative for benign) (Tavtigian et al., 2018). Panel A of figure 2.3 displays results for the 29 calibration variants. We note one outlier: p.L52P, which had relatively modest observational evidence in favor of pathogenicity (Odds_Path of 23:1), had an M2H result indicative of benignity. We also note that the residue L52 is unusual in that two benign calibration variants also fall there, p.L52F and p.L52I.

**Fig. 2.3.**
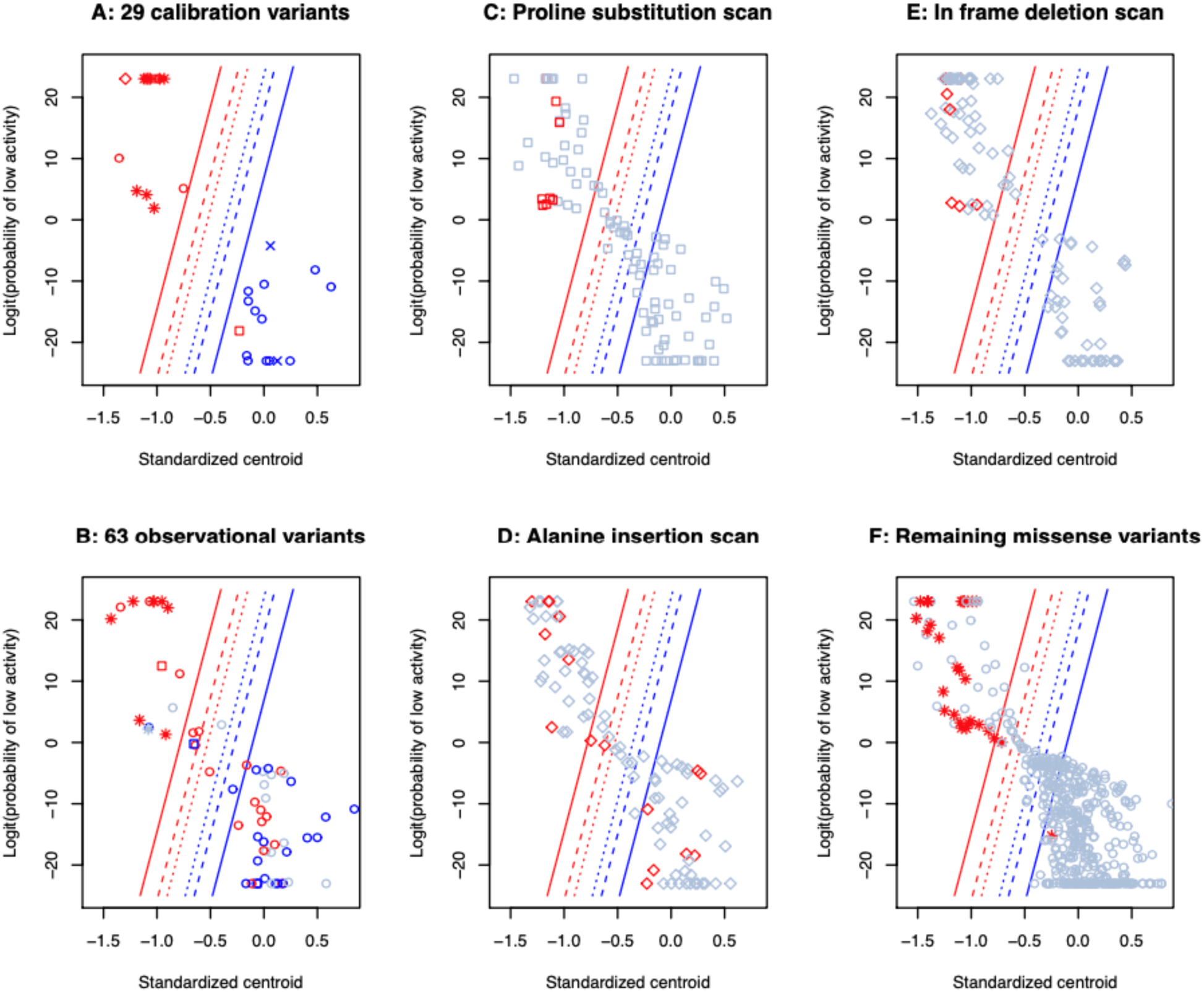
Distributions of all assayed sequence variants by standardized centroids (x-axes) and logit(probability of low activity) (y-axes). Odds of pathogenicity bounds (identical on all six panels): Red, evidence in favor of pathogenicity; Blue, evidence in favor of benign effect; Narrow dashed lines, ACMG supporting evidence i.e., odds = 2.08 or the inverse; Wide dashed lines, ACMG moderate evidence i.e., odds = 4.33 or the inverse; Solid liens, ACMG strong evidence i.e., odds = 18.7 or the inverse. **A**, The 29 sequence variants used for assay calibration. Red, observational evidence in favor of pathogenicity; Blue, observational evidence in favor of benign effect; Star, missense substitution at a C3HC4 residue; Square, other missense substitution to proline; X, cross-species super tolerated substitution; Circle, other missense substitution; Diamond, in-frame indel. **B**, The 63 additional observational variants. Red, variant reported in case series only; Blue, variant reported in population series only; Grey, variant reported in both series. Shapes as in A. **C**, Proline insertion scan. Red, proline substitution at a C3HC4 residue; Grey, proline substitution at another residue. **D**, Alanine insertion scan. Red, alanine insertion immediately adjacent to a C3HC4 residue; Grey, alanine insertion immediately adjacent to other residues. **E**, In-frame deletion scan. Red, in-frame deletions of a C3HC4 residue; Grey, in-frame deletions of another residue. **F**, Remaining missense substitutions not plotted in panels A-C. Red, missense substitution at a C3HC4 residue; Grey, missense substitution at another residue.

### Validation of the calibration

Beyond the calibration variants, an additional 63 snMS are present in breast and ovarian cancer case data from Myriad Genetics and Ambry Genetics that we described previously (Easton et al., 2007; Tavtigian et al., 2008; Li et al., 2020), and/or gnomAD v2 (non-TCGA) plus gnomAD v3 population data (Karczewski et al., 2020). To be clear, these snMS are individually very rare in the general population; the most common of them, p.R7C, was observed eight times in the ∼190,000 gnomAD subjects. Because the M2H data are comprehensive, all of these snMS are present in the functional assay data and can therefore be used as a validation series. Results are displayed in panel B of figure 2.3. Validation series snMS mapping into the upper left of the figure, in the M2H-based Strong evidence of pathogenicity bin, are highly enriched for clinical observations, and the crude odds ratios for the set of 15 snMS in the ACMG Strong evidence of pathogenicity bin was 10.8 (P=7.8×10^-5^) (table 2.2, table S2.1). In contrast, snMS falling in the lower right of the figure, in the M2H-based Strong evidence of benignity bin, show a mix of population and clinical observations; indeed, the odds ratio for the set of 40 snMS falling in the ACMG strong evidence of benignity bin was 0.68 (P=0.07). Data in the intervening bins are very sparse. Odds ratios for the combined Moderate & Supporting pathogenic, Indeterminate, and Supporting & Moderate benign bins were 2.95, 2.95, and 0.98, respectively; confidence intervals were very wide for all three. A test for trend across the ordered set of ACMG bins defined by the contour lines was indicative of the expected ordered series of odds ratios (P_trend_=4.8×10^-6^).

**Table 2.2.** Validation by odds ratio estimation

¥ For calibration regressions, we actually employed ceiling and floor values of
| Table 2.2. Validation by odds ratio estimation |  |  |  |  |  |
| --- | --- | --- | --- | --- | --- |
|  |  | # subjects / carriers |  |  |  |
| Category | # snMS | Clinical | gnomAD | Odds Ratio | 95% CI |
| Non-carriers |  | 193,000 | 190,069 | NA |  |
| M2H assay categories † |  |  |  |  |  |
| Strong pathogenic | 15 | 33 | 3 | 10.83 | 3.32 - 35.3 |
| Mod & Supp pathogenic | 4 | 3 | 1 | 2.95 | 0.31 - 28.4 |
| Indeterminate | 2 | 3 | 1 | 2.95 | 0.31 - 28.4 |
| Supp & Mod benign | 2 | 2 | 2 | 0.98 | 0.14 - 6.99 |
| Strong benign | 40 | 35 | 51 | 0.68 | 0.44 - 1.04 |
| M2H intersect Viability ¥§ |  |  |  |  |  |
| M2H-path & V-abnormal | 18 | 35 | 4 | 8.62 | 3.06 - 24.2 |
| M2H-path & V-normal | 1 | 1 | 0 | – |  |
| M2H-indet & V-abnormal | 1 | 1 | 0 | – |  |
| M2H-indet & V-normal | 1 | 2 | 1 | 1.97 | 0.18 - 21.7 |
| M2H-benign & V-intermediate | 3 | 1 | 2 | 0.49 | 0.04 - 5.43 |
| M2H-benign & V-normal | 38 | 34 | 51 | 0.66 | 0.43 - 1.01 |

† Ordered logistic regression across the five categories of snMS carriers returned $P=1.90 \times 10^{-8}$
¥ Ordered logistic regression across the six categories of snMS carriers returned $P=1.67 \times 10^{-8}$
§ One of the 63 validation variants, p.S36Y, was missing from the viability assay.

### Structural observations

Results for all remaining sequence variants that we assessed are summarized in panels C-F of figure 2.3 and figure S2.1. In general, single amino acid in-frame deletions were more likely to interfere with RING heterodimerization than were in-frame insertions of an alanine. Substitutions to proline were more likely to interfere with heterodimerization than other snMS. The seven canonical cysteines and one canonical histidine residues of the C3HC4 RING motif were particularly sensitive. In greater detail:

- At the seven canonical cysteines and one canonical histidine residues of the C3HC4 RING motif, 47 of 49 (96%) possible snMS had ACMG Strong evidence of pathogenicity. The two exceptions were p.C44G (Moderate pathogenic) and p.H41D (Strong benign).
- The BRCA1 RING structure includes two alpha helices that form a hydrophobic interface with the corresponding helices in BARD1 (Brzovic, Rajagopal, Hoyt, King, & Klevit, 2001). In helical wheel representation, the nine <u>a</u> and <u>d</u> residues (V8, V11, I15, M18, L22, L82, L86, I89, and F93) were sensitive to two kinds of substitution. First, at seven of the nine, substitutions to proline were Strong pathogenic; however, only two of these (p.L82P and p.L86P) could be reached by a snMS. Second, except at V8 and I89, the <u>a</u> and <u>d</u> residues were generally sensitive to replacement with a hydrophilic or charged residue; in total, 16 of 52 (31%) possible snMS had evidence of pathogenicity (12 Strong, 3 Moderate, 1 Supporting). The final residue of helix 2, D96, was also very sensitive; six snMS had Strong and the seventh (p.D96H) had Supporting evidence of pathogenicity.
- Across the remaining 81 RING domain residues, 35 of 483 (7.2%) possible snMS had evidence of pathogenicity (24 Strong, 9 Moderate, 2 Supporting).

### Stability of the calibration contours and strength of evidence bins

Stability of the calibration was assessed by iteration of the calibration step with each of the 29 calibrant variants excluded individually (panels A and B of figure 2.4) and then all 406 combinations of two calibrants excluded (panel C of figure 2.4). There were no instances where a Supporting bound moved far enough that a sequence variant switched from the Supporting pathogenic to Supporting benign bin (or vice versa), nor were there any instances where a sequence variant moved by more than one bin.

**Fig. 2.4.**
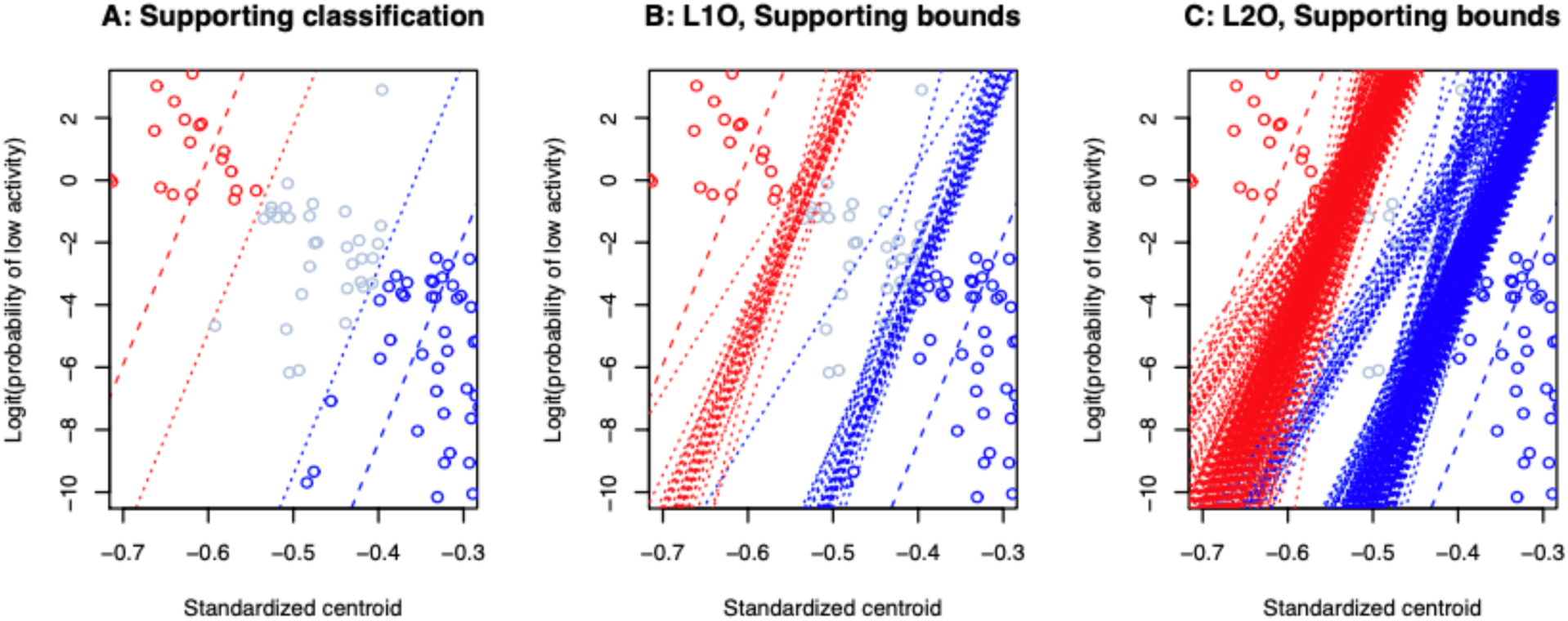
Leave one calibration variant out (L1O) and leave two calibration variants out (L2O) tests of sensitivity on location of the odds of pathogenicity bounds. **A**, Bounds obtained using the total calibration set (29 variants). Red, evidence in favor of pathogenicity; Blue, evidence in favor of benign effect; Narrow dashed lines, ACMG supporting evidence, i.e., odds = 2.08^±1^; Wide dashed lines, ACMG moderate evidence i.e., odds = 4.33^±1^. Also shown are the of variants with indeterminate or supporting evidence from the calibrated mammalian 2-hybrid assay. Grey circles display all assayed variants that fall between the supporting pathogenic and supporting benign bounds; these have indeterminate evidence, with categorical odds pathogenic of 1.0. Red circles falling between the supporting pathogenic and moderate pathogenic bounds display all variants with supporting evidence of pathogenicity, with categorical odds pathogenic of 2.08. Blue circles falling between the supporting benign and moderate benign bounds display all variants with supporting evidence of benign effect, with categorical odds pathogenic of 1/2.08. **B**, L1O supporting evidence bounds. As in A, but with the 29 supporting benign and supporting pathogenic bounds obtained from L1O recalculations superimposed. **C**, L2O supporting pathogenic evidence bounds. As in A, but with the 406 supporting pathogenic bounds obtained from L2O recalculations superimposed. **D**, L2O supporting benign evidence bounds. As in A, but with the 406 supporting benign bounds obtained from L2O recalculations superimposed.

### Cross analyses

The most comparable assay to the the M2H results described above is the nearly comprehensive cell viability assay reported by Findlay et al. (Findlay et al., 2018); intersection of the two assays includes 567 of the 591 possible RING domains snMS (96%) (with synonymous groups counted one snMS) (table 2.3). Of the 105 snMS for which the M2H assay finds evidence of pathogenicity (88 Strong, 13 Moderate, 4 Supporting), 87 had abnormal function in the viability assay, 8 had intermediate function, and 10 were considered functional. Thus we consider 87 to have concordant evidence of pathogenicity and 10 to be discordant. From the perspective of the viability assay, 116 were reported to have abnormal function. Of these, 87 had evidence of pathogenicity in the M2H assay, five were Indeterminate, and 24 had evidence of benignity. Thus we consider 87 to have concordant evidence of pathogenicity and 24 to be discordant.

**Table 2.3.** M2H versus viability assay confusion matrix.

| **Table 2.3. M2H versus viability assay confusion matrix.** |  |  |  |  |  |  |  |
| --- | --- | --- | --- | --- | --- | --- | --- |
|  |  | Viability assay categorizations |  |  |  |  |  |
|  |  | Abnormal |  | Intermediate |  | Normal |  |
| Row # | M2H assay categorizations | N | (C,V)† | N | (C,V)† | N | (C,V)† |
| 1 | Strong_path | 78 | (13,15) | 6 | (0,0) | 4 | (0,0) |
| 2 | Moderate_path | 6 | (0,3) | 1 | (0,0) | 6 | (0,1) |
| 3 | Supporting_path | 3 | (0,0) | 1 | (0,0) | 0 | (0,0) |
| 4 | Indeterminate | 5 | (0,1) | 4 | (0,0) | 8 | (0,1) |
| 5 | Supporting_benign | 1 | (0,0) | 2 | (0,0) | 13 | (0,0) |
| 6 | Moderate_benign | 5 | (0,0) | 8 | (0,0) | 30 | (0,1) |
| 7 | Strong_benign | 18 | (0,0) | 36 | (2,3) | 326 | (12,37) |
| 8 | M2H N/A & spliceogenic | 6 | (1,0) | 0 |  | 0 |  |

† (C,V) is (# snMS in the calibration series, # snMS in the validations series)
Note that two calibration variants, c.229\_237del (p.T77\_F79del) and c.271\_272TG>CA (p.C91H) were missing from the viability assay. However, neither is a snMS – they could only be included in the M2H assay because we specifically engineered them.
Note also that on validation variant, c.107C>A (p.S36Y) was missing from the viability assay.
Row 1 discordant snMS: p.L30W, p.K65R, p.F79S, p.F79L
Row 2 discordant snMS: p.I21F, p.I31F, p.I31S, p.L51F, p.I68T, p.Q81P
Row 3 discordant snMS: none
Row 4 discordant snMS: not applicable
Row 5 discordant snMS: p.P34R
Row 6 discordant snMS: p.A92V, p.A92T, p.G98V, p.L73R, p.E85G
Row 7 discordant snMS: p.V14G, p.V14F, p.I26T, p.V35A, p.H41D, p.K45I, p.L49P, p.L51P, p.P58H, p.N66Y, p.S72N, p.L73Q, p.S80I, p.L82F, p.E84K, p.I89N, p.Q94L
Row 8 spliceogenic snMS: c.211A>G (p.R71G), c.211A>T (p.R71W), c.212G>A (p.R71K), c.212G>C (p.R71T), c.212G>T (p.R71M), c.213G>C (p.R71S)

In addition, the viability assay identified six snMS at codon R71 with abnormal function because the underlying nucleotide substitution caused a splice defect. While these six snMS had normal function in the M2H assay, our prior sequence analysis predicted spliceogenicity (Vallee et al., 2016), which means *a priori* that the M2H assay cannot be used as evidence of benignity for these unless the variants have been found to have normal activity in a mRNA splicing assay, which is not the case here. Therefore, we do not consider these as false benign results from the M2H assay, and we also note that there were zero nucleotide substitutions with an RNA defect in the viability assay that were not predicted beforehand by our sequence analysis.

Intersection between the M2H and viability assay results provides one more route to validating the Odds_Path assigned above. Among the 87 snMS with concordant evidence in favor of pathogenicity, 13 were calibration snMS. Because these were by definition observed in a case before study inception, they have to be excluded from the analysis that follows. Of the remaining 87-13=74 concordant snMS, 16 were observed in one or more of the cases included in the validation series. If the discordant snMS are actually pathogenic, a similar proportion (16/74) should be observed in a validation series case. Thus of the 10 discordant snMS with evidence of pathogenicity coming from the M2H assay, we should expect to have observed (16/74)×10=2.2 in a validation series case and observed one. The discordant snMS observed was p.I68T. This snMS was observed in one case from Ambry, falls at a residue where two other snMS had concordant evidence of pathogenicity (p.I68K and p.I68R), had moderate evidence of pathogenicity in the M2H assay, and was reported to have normal function in the viability assay. Of the 24 discordant snMS with evidence of pathogenicity coming from the viability assay, we should expect to have observed (16/74)x24=5.2 in a validation series case and found zero.

Therefore, arguing from anecdotal evidence and lack of evidence, we estimate that about ½ of the snMS with a pathogenic result from the M2H assay but intermediate or normal function in the viability assay, have a false pathogenic M2H result, resulting in a proportion pathogenic of (96/105)=0.914. Starting from the estimated proportion pathogenic among BRCA1/2 key domain missense substitutions and in-frame indels, 0.35 (Easton et al., 2007; Li et al., 2020), application of Bayes’ rule reveals that a pathogenic result from the M2H assay is associated with Odds_Path of 19.8:1 – meeting the criterion for ACMG Strong evidence of pathogenicity (note also that if the analysis were focused on the M2H Strong pathogenic group, the Odds_Path would be 32:1). If we estimate that about 1/3 of the snMS with an abnormal function result from the viability assay but Indeterminate or Benign M2H result, Bayes’ rule reveals that an abnormal function result from the viability assay is associated with Odds_Path of 9.5:1 – meeting the criterion for ACMG Moderate+ evidence of pathogenicity.

### Variant classification model

The ACMG-compatible Bayesian points system (Tavtigian et al., 2018; Tavtigian et al., 2020) can be used to integrate several kinds of data toward classification of RING domain snMS, employing the following codes and logic. We present some of these ACMG evidence criteria in pairs when they are mutually exclusive, starting with information about the position of the snMS, then effect of the alternate amino acid, and finally population and case observational data. The human subjects observational data are added last because these data are most likely to evolve with addition of clinical information.

PM1: “…well-established functional domain (e.g., active site of an enzyme) without benign variation”. We note here that the eight canonical C3HC4 residues explicitly meet this criterion. But the <u>a</u> and <u>d</u> positions of the two interface alpha helices don’t quite meet this criterion because we can see from our reference BRCA1 protein multiple sequence alignment (Tavtigian et al., 2008) that a number of alternate amino acids are within the evolutionarily tolerated range of variation (e.g., V11A, V11I, I15L, M18L…).

Therefore, for these key <u>a</u> and <u>d</u> residues, we deprecate to PM1_Supporting. Moreover, at the remaining 81 RING domain residues, several snMS have been classified benign, so we deprecate these residues to PM1_Indeterminate. (+2, +1, or 0 points)

PM5: “Novel missense change at an amino acid residue where a different missense change determined to be pathogenic has been seen before”. (+2 points)

PP3 vs BP4: “Computational tool”. Align-GVGD is well-attested for use with BRCA1 RING and BRCT domain misssense substitutions. Here, we use a conservative implementation with C0 assigned -1, C15-C35 assigned 0 and C65 assigned +1 point.

PS3 vs BS3 “Well-established *in vitro* or *in vivo* functional studies….” As calibrated above, M2H scores range from +4 to -4 points. Viability assay results are assigned +3, 0, or -3 points. Total scores are limited to +/-4 pending community consensus.

PM2_Supporting vs BS1_Moderate: “absent from controls….” (the hereditary breast and ovarian cancer VCEP has reduced PM2 to Supporting, and the HBOC VCEP applies BS1_Moderate to VUS with more than two observations in gnomAD) (Parsons et al., 2019). Thus we also have PM2_BS1_Indeterminate if a VUS has one or two observations in gnomAD. (+1, 0, or -2 points).

To avoid circularities, the calibrated functional assay results should not be applied to the calibration series variants summarized in table 2.1. However, the calibrated functional assays and points-based evidence integration can be applied to the 63 validation variants. The resulting classifications are depicted in figure 2.5 and summarized in table S2.1. Receiver operating characteristic (ROC) area analysis revealed that the ROC area of the classification model increased sequentially with addition of each evidence criterion. However, it should be acknowledged that the final addition (PM2 and BS1) is directly dependent on case and control observational data and therefore circular in a ROC analysis that uses case/control status as the indicator variable.

**Fig. 2.5.**
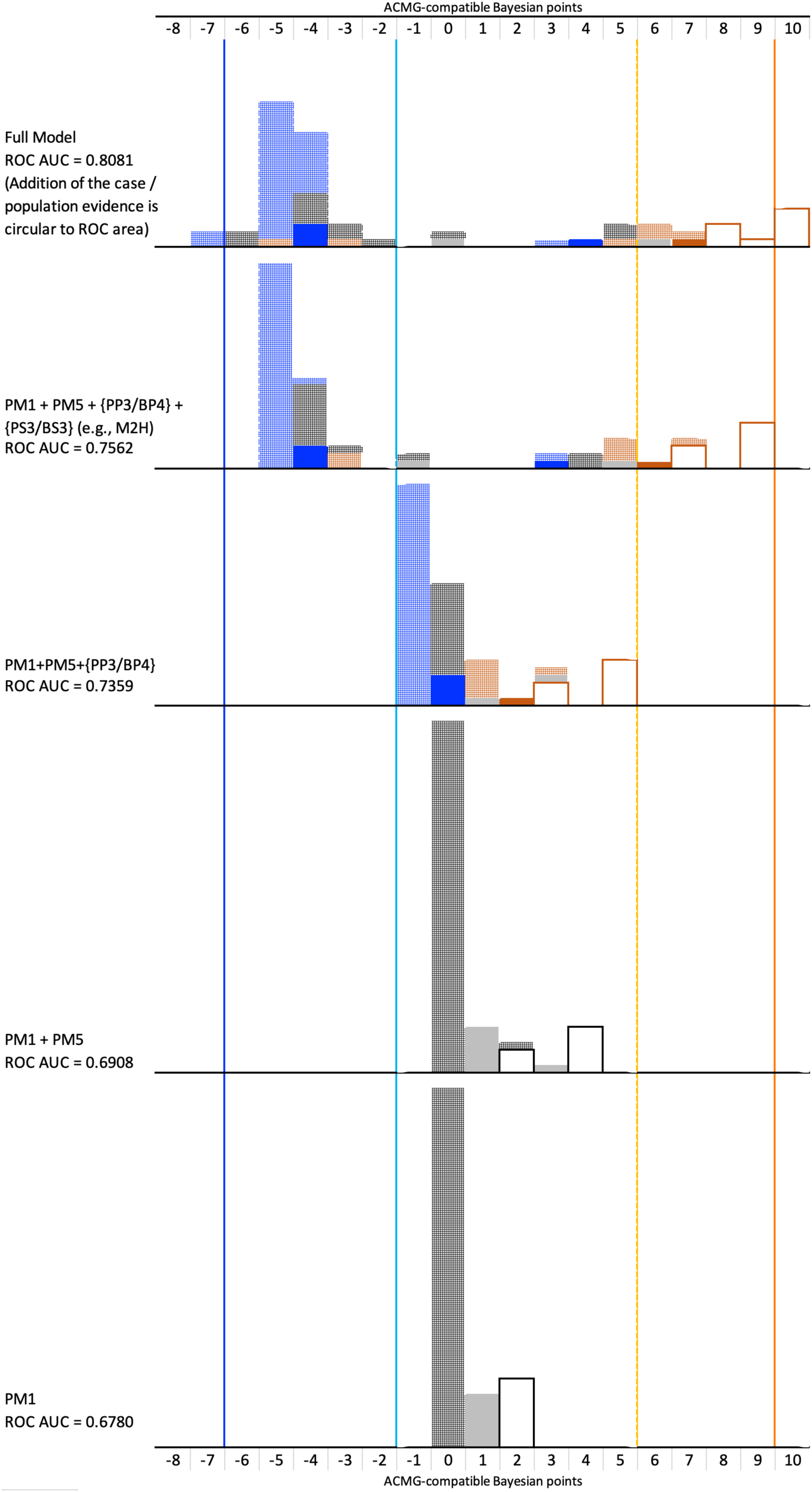
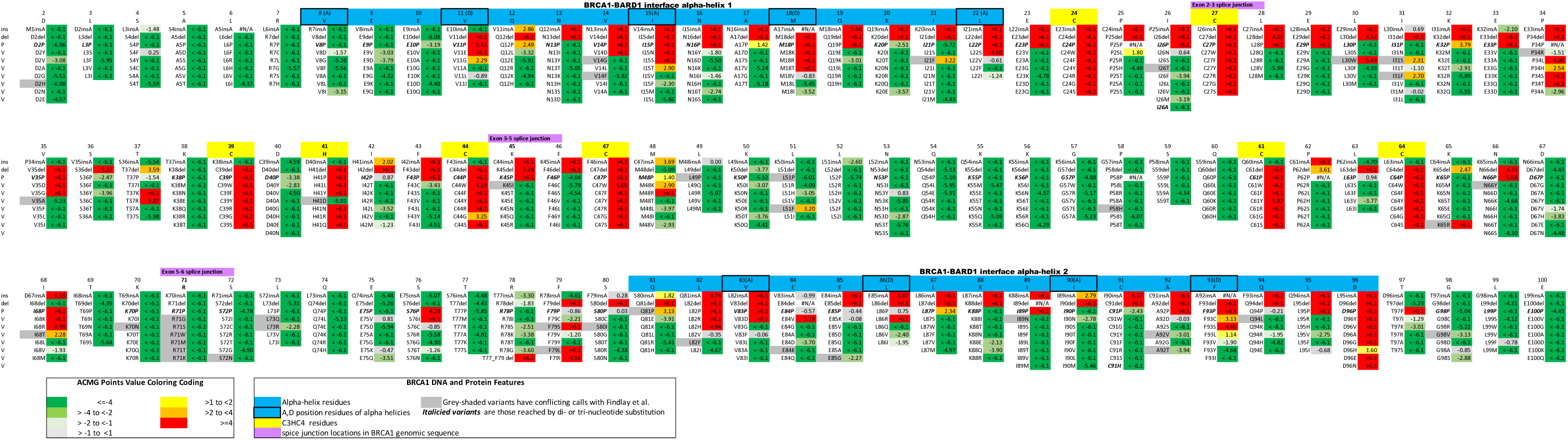
Sequential addition of evidence criteria to the missense substitution classification model, as applied to the 63 validation series missense substitutions. The sequence builds from the bottom of the figure up. Model components and ROC AUC are given at the left. Vertical blue line, Benign threshold. Vertical cyan line, Likely benign threshold (ACMG rule LB-II). Vertical gold line, Likely pathogenic threshold. Vertical orange line, Pathogenic threshold. Solid border, no-fill boxes, C3HC4 residues. Solid fill, alpha helix a,d residues. Hatched fill, all other RING domain residues. Deep orange border or fill, Align-GVGD C65 missense substitutions. Grey fill, Align GVGD C15-C55 missense substitutions. Blue fill, Align-GVGD C0 substitutions.

Overall, five of these snMS obtain scores of 10 (Pathogenic); these all fell at C3HC4 residues and had concordant functional assay evidence of pathogenicity.

Nine obtained scores of 6 to 9 (Likely Pathogenic). These fell at all three types of RING domain positions, including two (p.L28P and p.P34S) at non-C3HC4, non alpha-<u>a</u>, <u>d</u> residues not harboring an already known pathogenic snMS. Nonetheless, the nine Likely Pathogenic variants all had concordant functional assay evidence of pathogenicity.

Seven obtained scores of -1 to 5 (VUS). These had very mixed profiles with, for instance, functional assay results ranging from concordant evidence of pathogenicity to concordant evidence of benignity. One of the more interesting results was p.L22F. This substitution fell at an alpha-<u>a</u>, <u>d</u> residue harboring a known pathogenic substitution (p.L22S); nonetheless, the functional assay results provided concordant evidence of benignity, and the final score was 0.

Forty obtained scores of -2 to -6 (Likely Benign). These again had mixed profiles, but 37 fell at non-C3HC4, non alpha-<u>a</u>, <u>d</u> residues. 35 had concordant functional assay evidence of benignity, and the remaining five had evidence of benignity from one assay and indeterminate evidence from the other.

The remaining two snMS had scores of -7 (Benign). They were both physicochemically conservative substitutions (p.D2N and p.Q80R) falling at non-C3HC4, non alpha-<u>a</u>, <u>d</u> residues and had concordant assay-based evidence of benignity.

We also note that all nine snMS falling at C3HC4 residues were classified either Likely Pathogenic or Pathogenic. In contrast, of the seven snMS falling at alpha-<u>a</u>, <u>d</u> residues, two were classified LP, two VUS, and three LB. Overall, the proposed model provided clinically useful classifications for 56 of the 63 (89%). LB was by far the most common result (40 of 63), and LP was the next most common result (9 of 63).

## DISCUSSION

### Assay development and coverage

Previously, we adapted a chemiluminescent Mammalian 2-hybrid (M2H) assay (Paquette et al., 2018), to assess BRCA1 RING domain missense substitutions and in-frame indels. Here, the chemiluminescent assay was upgraded to four-color fluorescent protein assay that allowed gating on expression of both proteins required for the two-hybrid assay and flow sorting based on expression from two M2H reporter loci. Although the assay relied on transfection of BRCA1 and BARD1 expression constructs into individual wells of a multi-well plate and was thus moderate-throughput, it could be read out by flow sorting, massively parallel sequencing, and sequence analysis much as higher throughput multiplex assays of variant effect (MAVEs) enabling analysis of 853 sequence variants. In principal, the assay could be upgraded to high-throughput by modifying the reporter cell line to contain a target insertion site for the mutagenized expression construct or by placing the expression construct, re-designed for CRISPR mutagenesis, into a genomic context. The assay design included silent barcodes within the BRCA1 M2H expression construct, allowing positive detection of three or more independent replicates of each desired RING domain variant, and the assay was driven to 100% completion.

### Calibration and Validation

The output of the M2H assay was calibrated as Odds_Path and then reduced to quantitatively defined ACMG strength of evidence categories (Richards et al., 2015; Tavtigian et al., 2018) and ACMG-scaled points (Tavtigian et al., 2020; Garrett et al., 2020). This formulation allows direct combination of the functional assay results with other data towards BRCA1 RING domain variant classification, so long as the calibration is reasonably accurate. To that end, after the M2H assay output was calibrated using patient observational evidence for or against pathogenicity from a set of 28 missense substitutions plus one IFD, we made two independent tests of the calibration.

First, based on mutation screening data from almost 200,000 breast and ovarian cancer cases plus a similarly sized population sample, 63 additional snMS were observed one or more times. The odds ratio associated with the Strong pathogenic bin, 10.8, was consistent with a grouping that is highly enriched in pathogenic variants. The odds ratios associated with the Moderate and Supporting pathogenic bins, and the Indeterminate bin, 2.95 (with very wide confidence intervals), are consistent with either enrichment for pathogenic variants or else a high proportion of moderate-risk variants. The odds ratios associated with the Supporting and Moderate benign bins, and the Strong benign bin, 0.98 and 0.68, respectively, are indicative of groupings consisting almost entirely of benign variants. Moreover, the odds ratio trend across the strength of evidence bins is consistent with the idea that stratification by calibrated functional assay result corresponds to stratification by cancer risk.
Second, because the M2H results are comprehensive and the Findlay et al viability assay results nearly so, cross-comparison of the results gave an almost comprehensive view of the discordance rate between the two assays (table 2.3). Under the assumption that the discordance rate provides an upper estimate of the error rate, comparison of the discordance rate to the proportion pathogenic expected in each category is informative. Of 88 snMS in the M2H Strong pathogenic bin, 47 fell at a C3HC4 residue and none of these had a discordant result. An additional 18 fell at an alpha-<u>a</u>, <u>d</u> residue and none of these had a discordant result. The remaining 23, with four discordant results, fell at PM1_Indeterminate positions. For these, the minimum expected proportion pathogenic is the Bayesian combination of the global ACMG Prior_P of 0.102, PM1_Indeterminate Odds_Path of 1:1, and PS3 Odds Path of 18.7:1, and thus a Post_P of 0.68. As 19/23>0.68, the expectation was met. Thus, from this analytic point of view, calibration of the M2H assay also met or exceeded expectations for the ACMG PS3 category irrespective of the actual pathogenicity of the discordant variants.
Of 380 snMS in the M2H Strong benign category, one fell at a C3HC4 residue (p.H41D) and was discordant. An additional 24 fell at alpha-<u>a</u>, <u>d</u> positions, and two were discordant. The remaining 355 fell at PM1_Indeterminate positions, and 15 were discordant. Across these three PM1 groups of amino acid positions, the weighted average Odds_Path was 1.077:1. Hence the maximum expected proportion pathogenic is the Bayesian combination of the global ACMG Prior_P of 0.102, weighted average PM1 Odds_Path of 1.077:1, and BS3 Odds Path of 0.0534:1, and thus a Post_P of 0.0065. That calculation implies that there should be at most two or three pathogenic snMS among the 380 snMS with M2H strong benign results. Thus either only a small fraction of the discordant snMS in this group are pathogenic, or else the assay does not actually meet the ACMG BS3 criterion. Here, we note two points: (1) If the proportion pathogenic among the 18 discordant snMS were similar to those with a concordant pathogenic result, we would expect to have observed 4 in the validation series – yet none wore observed. (2) Even if nine of these 18 discordant snMS are actually pathogenic (potentially, an over-estimate), then the proportion pathogenic in the M2H Strong benign category would be 9/380=0.024 – which is below both the ACMG and IARC thresholds for classification as LB.

Thus we are confident that an M2H Strong pathogenic result meets the ACMG PS3 criterion, but perhaps counter-intuitively cannot be sure that an M2H Strong benign result meets the ACMG BS3 criterion. Furthermore, there are simply not enough patient or population observational data to test whether combined results from the M2H and viability assays provide more than ACMG Strong pathogenic or benign evidence. For this reason, we recommend the combined results of the two assays should be limited to Strong pathogenic and Strong benign (+/- 4 points) until the question can be evaluated on larger datasets.

### Limitations and Sources of Error

One may envision that a functional assay should yield perfect results; nonetheless, “Strong” odds of 18.7:1 in fact allow for an appreciable error rate. As the RING M2H experiment involves a multiple test of 604 for the comprehensive set of snMS, and almost 900 when all variant types are included, some stochastic error is expected. Technical errors are also possible. For example, outside of the moderate throughput workflow, we performed 1-by-1 assays on a small number of variants that had unexpected results (not shown). Among these, p.K65R was a unique outlier because it reproducibly showed reduced function in the parallel assay but had aberrant BFP:YFP expression when assayed alone, potentially indicative of interference between the missense substitution and the P2A cleavage signal in its expression construct. Mechanistic error would be possible if there can be separation of function between BRAC1-BARD1 RING domain heterodimerization and another function that is critical to tumor suppression. Here, we note that a reciprocal question applies to other assays: if the M2H-autoubiquitination separation of function variant p.I26A has not been tested in other assays, there is concern that they could produce false pathogenic results to the extent that their readout is dependent on, or susceptible to, ubiquitination activity. In addition, cDNA based assays are liable to mechanistic errors because they are insensitive to mRNA splicing defects. Here, we note that our conception of PP3/BP4 computational evidence and PS3/BS3 functional assay evidence involves separate tracks for snMS severity and damage to mRNA splicing. Within these evidence codes, a computational prediction for a splice defect can only be negated by functional assay evidence against a splice defect, which the cDNA-based M2H assay cannot provide. However, the genome edited viability assay can provide this evidence (Findlay et al., 2018), which is an important complementarity between the two assays.

In addition, there is concern about the analytic status of moderate risk variants, such as the *BRCA1* BRCT domain snMS p.R1699Q (Spurdle et al., 2012). Conferring ORs between 2 and 4-5 (Easton et al., 2015), there are not as yet any clear examples from the RING domain, so it is not clear how much activity these would have in the M2H assay. Based on sparse data, the validation series (table 2.2) ORs associated with the M2H Moderate and Supporting pathogenic bins, and the Indeterminate bin, are consistent with the presence of moderate-risk variants. Under a scenario where most of the evidence towards classification of very rare snMS comes from a combination of functional assay and computational data, plus exceedingly sparse human observational data, one might guess that most genuinely moderate-risk snMS will rest as VUS, but the issue poses an analytically difficult question.

### Classification Model

The classification model proposed here is the points system that is based on the quantitative Bayesian interpretation of the qualitative ACMG variant classification guidelines. We employed five ACMG evidence criteria that are universally available for observed snMS: PM1 (functional domain) with added analytic precision to allow +2, +1, or 0 points; PM5 (known pathogenic missense substitutions at this residue), +2 or 0 points; the pair PM2_supporting and BS1_moderate, which are related to the number of observations in the case and population data sets and allowed values of +1, 0, or -2 points; and pair PP3 and BP4 (computational tool), supplied by Align-GVGD and allowed +1, 0, or -1 points. Finally, PS3 and BS3 conveys the functional assay results and was allowed values from +4 through 0 to -4. Noting that addition and subtraction of points corresponds to multiplication of the associated Odds_Path, we combined results from the comprehensive M2H and viability assays by adding their scores for each individual variant while applying limits of +4 and -4 to those sums. Because the viability assay appears to have a higher false pathogenic rate than the M2H assay, we allowed scores of +3, 0, and -3, and note that deriving in-between scores such as +2 and +1 would require a complete re-analysis of the underlying viability assay data. Overall, the approach was very efficient, with 89% of observed variants (i.e., the table S2.1 validation series variants) reaching a clinically applicable category. While the classification could also be applied to unobserved snMS, we hesitate to do so because the affectation status of the first few observed carriers could change the result; should it be decided that unobserved snMS can be classified, we note that the point totals available from this system can reach LB or LP, but cannot reach either the B or P thresholds.

### Summary

Applied to human disease susceptibility genes, multiplex assays of variant effect can be exceptionally efficient source of data towards variant classification. But they often rely on a cell viability phenotype that may be associated with but not necessarily causally connected to the mechanism underlying susceptibility. Here, we have driven a protein interaction assay that is mechanistically central to the function of the BRCA1-BARD1 heterodimer to completion. Calibration followed by two approaches to validation demonstrate unambiguously that loss of function in the M2H assay provides ACMG Strong evidence in favor of pathogenicity. Calibration of the assay results as Odds_Path allows results from each individual snMS to feed directly into a simple points-based variant classification system. Yet, despite its simplicity, the system is sufficiently effective that 89% of the validation series snMS could be classified as either P/LP or LB/B. In one of the first *BRCA1* gene discovery confirmation papers, Castilla *et al* opined that “False positives (missense mutations which are later revealed to be rare benign polymorphisms) will be a potentially serious problem until functional assays for *BRCA1* can be established” (Castilla et al., 1994). And so it is. Among the 27 calibration series variants that are observed snMS, 15 (56%) fell in the M2H Strong evidence of pathogenicity bin. Among the 56 validation series snMS classified as either P/LP or LB/B, only 14 (25%) fell into the P or LP category. Finally, among the 370 snMS with concordant results in the M2H and viability assays that were neither part of the calibration nor validation series (thus not observed among ∼400,000 subjects and therefore exceedingly rare), 56 (15%) had functional assay evidence of pathogenicity. Thus an important effect of the functional assays, coupled to points-based classification, is to improve the efficiency of recognizing benign snMS among the RING domain snMS. Still, several issues remain to be resolved, including: resolving the status of snMS with discordant results between the M2H and viability assays; clarifying expected results and appropriate scoring for moderate-risk snMS; systematic inclusion plus implications towards assay calibration of separation-of-function snMS, and defining the scoring range applicable to combined results from multiple functional assays, taking into account that they may not be mechanistically independent of each other. Nevertheless, among the 56 rMS with concordant functional assay evidence in favor of pathogenicity, 47 also have computational evidence of pathogenicity and another five are substitutions to proline that fall within one of the two BRCA1-BARD1 interface alpha helices; these are particularly likely to be classified as LP once human observational data become available.

## METHODS

### Pre-set analysis choices

A number of key analysis parameters were set before the resulting analyses were conducted. These included:

1. Calibration would be based on some sort of multivariate logistic regression using human observational data, and excluding computational or functional assay data.
2. The calibration set of RING domain sequence variants used inclusion thresholds of human observational data having 0.10 < Post_P < 0.90, starting from an assumed BRCA1/2 key domain Prior_P of 0.35 (Easton et al., 2007; Li et al., 2020).
3. Because the calibration set had more pathogenic than benign calibrant variants, it was decided in advance to include the two protein multiple sequence alignment based variants p.K45R and p.C91H (Paquette et al., 2018).
4. The calibration would be subject to an independent validation using observational data and an odds ratio approach.
5. For the missense substitution classification model, the order of addition of the individual evidence criteria was set before the sequential ROC AUCs were estimated.

### Human subjects information

Human subjects data were gathered from several published studies or data tables underlying those studies. Some *BRCA1* mutation screening data are from 68,000 full-sequence *BRCA1/2* tests carried out at Myriad Genetic Laboratories before 2008. For these, a test request form must have been completed by the ordering health care provider, and the form must have been signed by an appropriate individual indicating that ‘‘informed consent has been signed and is on file.’’(Easton et al., 2007; Tavtigian et al., 2008). Additional *BRCA1* mutation screening data are from 138,342 multigene panel tests performed by Ambry Genetics and “exempted from review by the Western Institutional Review Board.” (Li et al., 2020). Odds_Path derived from personal and family cancer history and segregation were gathered from the BRCA1/2 “Ex-UV database”, the ENIGMA consortium, and the more recent Ambry Genetics study (Vallee et al., 2012; Parsons et al., 2019; Li et al., 2020). Population mutation screening data were obtained from non-TCGA gnomAD v2 plus non-overlapping sequences from gnomAD v3 (Karczewski et al., 2020).

### Assay

The basic dual reporter M2H assay used here was described previously described in a low throughput format (Paquette et al., 2018). Three upgrades were made to create a moderate throughput format suitable for comprehensive assay of the *BRCA1* RING domain. First, we had noted from RT-PCR of the transfected BRCA1-GAL4 and BARD1-VP16 expression constructs that the lowest flow sorted M2H activity bin (panel C of figure 2.1) had relatively low expression of BARD1-VP16. This was rectified by modifying the BARD1-VP16 construct to include yellow fluorescent protein preceded by a P2A cleavage sequence, and then gating transfected cells for expression of both blue and yellow fluorescent protein. Second, the repertoire of silent barcodes was expanded to allow for eight distinct barcodes within each of the three 33-amino acid BRCA1 segments that were array synthesized. Finally, to support higher throughput, plasmid minipreps and transfection of HEK293 cells were conducted in 96-well format. To avoid edge effects during tissue culture, the outer rows and columns of the 96-well tissue culture plates were not transfected, i.e., transfections used 60 wells per plate.

### Statistical methods

The data were read in from two tables. The first was a small table giving the total cell counts for in each bin for each experiment. We denote as *N_j,l_* the cell count in bin *l* for experiment *j* with *j* ɛ {1, . . . 7} and l ɛ {1, . . . 6}. The second was a large table containing all the sequence counts of each variant in each bin, on each barcode, for each experiment. Denote by *Y_i,j,k,l_* the sequence count for variant *i* for experiment *j* on barcode *k* in bin *l*. Note that due to PCR amplification preceding sequencing, the sequence counts in any bin for any experiment are far larger than the corresponding cell counts. Note also that the cell counts are not random but an experimental design choice.

All data manipulation, analyses and statistical graph drawing were done using the R statistical environment (R Core Team 2015).

### Centroids

The assay was designed so that the bin indexes correspond roughly to distance from the origin in 2-dimensional red-green fluorescence activity space. We use this to define a *centroid* as an approximate measure of centrality of the distribution of cells in this red-green space, as follows. We use the conventional dot notation to indicate summation over an index so that

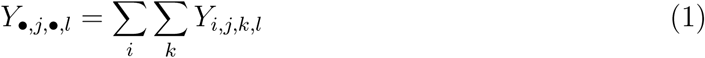

is the total sequence count in bin *l* for for experiment *j*. If *X_i,j,k_* is the centroid for the distribution of cells of variant *i* in experiment *j* on barcode *k*, then,

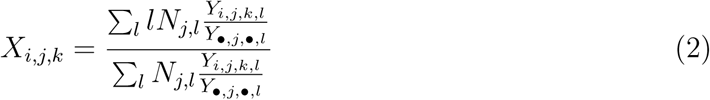

To test for and estimate experiment and barcode effects on the centroid we used standard linear regression methods to fit the model

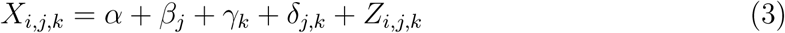

where *α* is the intercept term, */3_j_*and *’Y_k_*are the main effects for experiment *j* and barcode *k*, *δ _j,k_* is the experiment by barcode interaction term, and each *Z_i,j,k_* is an independent random Normal(0*, σ* ^2^) error. We found strong evidence in favor of including all the explanatory variables in equation (3). A simple plot of the residuals from this fit, panel C of figure 2.2, showed clear evidence for bimodality in the data not explained by this model.

### Fitting a bimodal model using the EM algorithm

In order to fit a bimodal model to the centroid data, we let *Q_i_* be an indicator variable for each variant *i*, with 0 indicating that the variant has activity levels similar to the wild type, and 1 indicating low activity level. Clearly, these indicators are not observable, however, we exploited the bimodality in the data to estimate them using the expectation-maximization, or EM, algorithm (Demster et al., 1977). This is a standard approach to estimating such hidden variables. We do this by applying standard weighted regression methods in an iterative fashion.

At any stage in the iterative process, let *P_i_* be the expected value of *Q_i_*, or equivalently, the probability that *Q_i_* is 1. We initialize the process by setting *P_i_* to 0 when *i* is the wild type, and 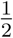 for all other variants.

We then considered the linear regression

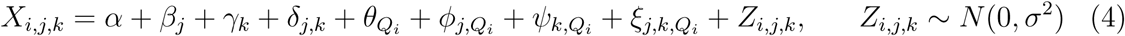

that is, to the model in equation (3) we added a main effect for the activity indicator and all terms for interactions with experiment and barcode. For any given set of *{P_i_}* we can fit this model using standard weighted regression where we included each observation *X_i,j,k_* twice: once with *Q_i_* set to 1 with weight *P_i_* and once with *Q_i_* set to 0 with weight 1 *P_i_*, so that the total weight for each observation is 1. This is the M-step of the EM algorithm and yields maximum likelihood estimates for the parameters of the model.

Given the set of parameter estimates for model (4) we can obtain *µ_i,j,k,_*_0_ and *µ_i,j,k,_*_1_ the expected values of *X_i,j,k_* under the hypotheses that it has wild type and low activity respectively. We also obtain an estimate of *17*^2^ the variance, which is assumed to be the same under each hypothesis. Thus, if variant *i* has wild type activity its distribution is *N* (*µ_i,j,k,_*_0_*, σ* ^2^) and is *N* (*µ_i,j,k,_*_1_*, σ*^2^) if it has low activity. We can then update our estimates of *{P_i_}* using

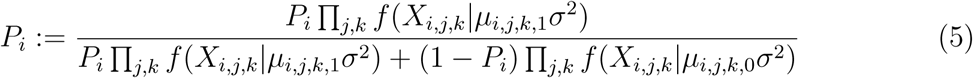

where *f* (*x|µ σ*^2^) is the density function for a *N* (*µ, σ*^2^) random variable. This is the E-step of the EM algorithm and gives us new estimates for the expected values of *Q_i_*.

Using the initial values given above, we iterated through the M-step and E-step 50 times. This was more than sufficient to achieve convergence which we observed at about 20 iterations.

Thus, for each variant, the EM-algorithm yields an estimate of *P_i_* the probability that it has low activity, (*PLA*). In addition we define a *standardized centroid* to give an overall estimate of its activity level. We denote this as *C_i_* and define it as the mean of a variant’s observed centroids with each corrected and scaled to account for experiment and barcode effects.

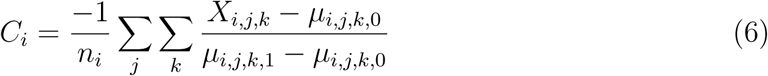

where *n_i_* is the total number of observations for variant *i*.

### Calibration to prior evidence for pathogenicity

From previous work by other authors we have 15 variants with strong evidence of pathogenicity and 14 with strong evidence of benignity, for a total of *n* = 29 variants that we can use for calibration. This evidence is expressed as the odds in favor of pathogenicity derived from the previous assay or analysis. In order to avoid extreme values we truncated the odds to be between 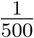 and 500. We assumed that a randomly chosen variant has a prior probability *π* = 0.35 of being pathogenic (Easton et al., 2017; Li et al., 2020), and, using Bayes rule, we combined this with the observed odds to obtain *q_i_*, the probability that each of the calibration variants is pathogenic.

In order to characterize the results of our assay for pathogenic and benign variants, we assumed that the logit of the PLA and the standardized centroid had a bivariate Normal distribution with different mean vectors depending on pathogenicity but the same variance-covariance matrix under both hypotheses. We let 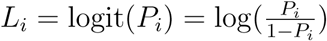.

The parameters for these bivariate Normals are (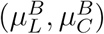) the mean vector for benign variants, (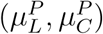) the mean vector for pathogenic variants, 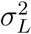 the variance of logit(PLA), 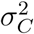 the variance of standardized centroid, and *⇢* the correlation coefficient for logit(PLA) and standardized centroid. The variances and covariances are the same for all variants. These are estimated using standard weighted estimation as follows

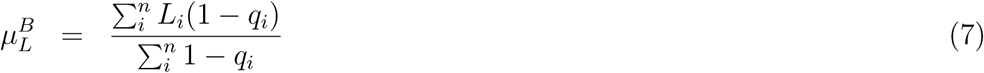

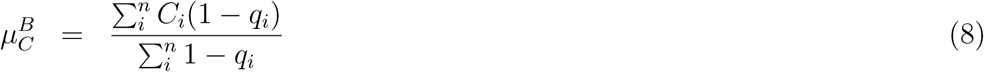

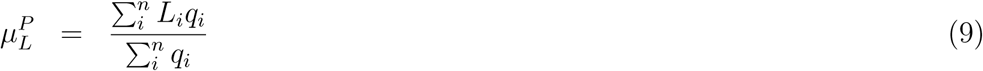

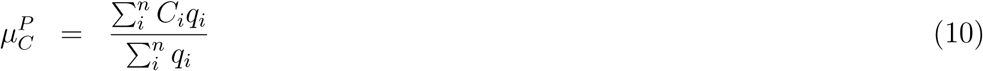

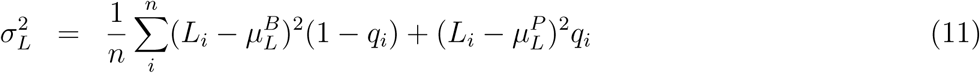

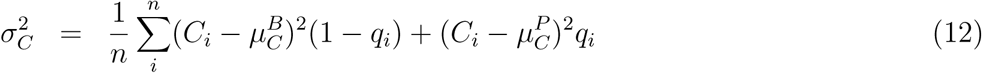

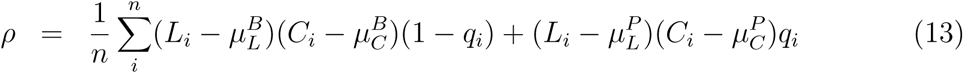

We can now use these estimates to obtain the evidence in favor of pathogenicity for any variant in our assay. Assuming that the (non-calibrant) variant has *L* = logit(PLA) and *C* = standardized centroid, we express this evidence as the log of the ratio of the probability of these observations under the hypothesis of pathogenicity over their probability under the hypothesis of benignity:

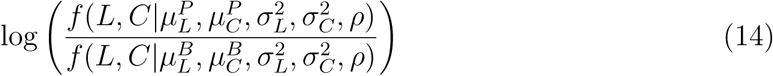

where *f* () here is the bivariate Normal density function. This can be obtained by the following straightforward calculations:

• Transform *L* into a standard Normal under each hypothesis to get

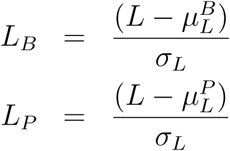

• Transform *C* into a standard Normal under each hypothesis to get

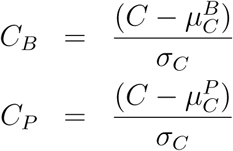

• Then the log likelihood ratio in favor of pathogenicity is given by

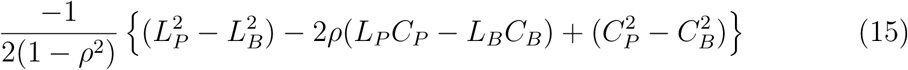

We, thus, computed the log likelihood ratio in favor of pathogenicity for all variants in the assay. Figure 2.3 displays scatter plots of (*L_i_, C_i_*) for each variant and superimposed critical contours of the log likelihood surface.

In order to investigate the sensitivity of the calibration to the available variants we made 29 leave-one-out (L1O) repeats of the above analysis leaving out every calibration variant in turn. Then, in a leave-two-out (L2O) run we made 406 replicates leaving out in turn every pair of observations. For each replicate we calculated and plotted the new critical contours of the log likelihood surface; these are displayed in figure 2.4.

Finally, frequentist odds ratios and ROC AUCs were estimated in Stata 15.1 (StataCorp).

**Table S2.1.**
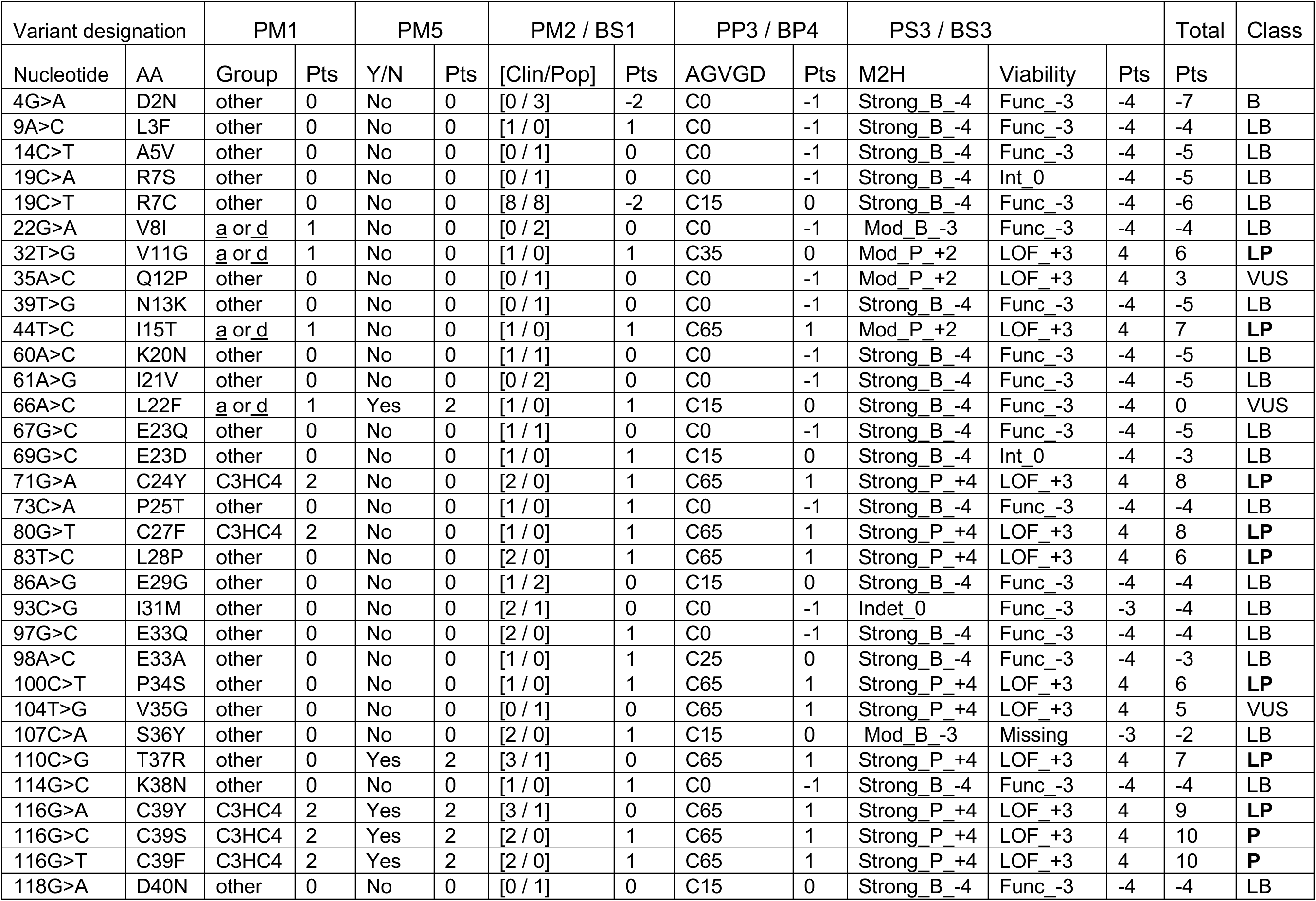
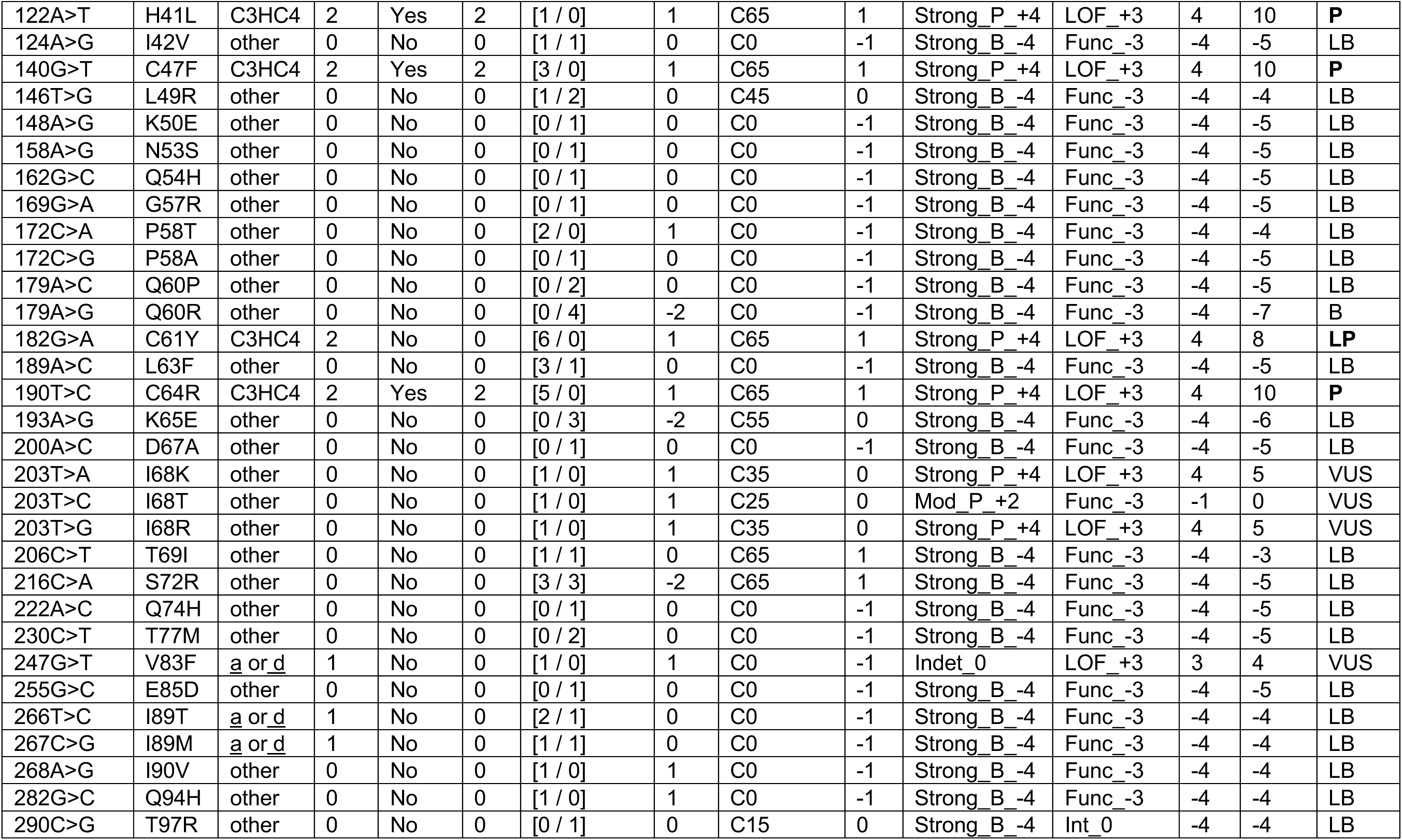
Points-based classification applied to the 63 validation missense substitutions.

## ACKNOWLEDGEMENTS

This work was supported by the United States National Institutes of Health (NIH) grants R01CA164944, R01CA121245, P30CA042014, S10RR026802, and the Canadian PERSPECTIVE I&I Project through the Canadian Institutes of Health Research (GP1-155865). BAT was an Australian National Health and Medical Research Council CJ Martin Early Career Fellow. We also thank the University of Utah Mutation Generation and Detection Core.

